# Engineered proteins to restore and enhance neurotransmitter release

**DOI:** 10.64898/2026.10.23.761409

**Authors:** Morven Chin, Ke Yang, Ricardo C. López, Chao Tan, Pascal S. Kaeser

**Author notes:** equal contribution (Morven Chin, Ke Yang, Ricardo C. López, Chao Tan).

## Abstract

Synapse dysfunction is a hallmark feature of many neurological diseases. Synaptic transmission is initiated in nerve terminals, where active zone protein complexes control vesicle fusion for neurotransmitter release. Here, we engineer proteins that mediate active zone functions and demonstrate their use for reconstituting and strengthening neurotransmission. A protein composed of Ca_V_β4 and the RIM1 zinc finger, β4-Zn, restored neurotransmitter release after active zone disruption through recruitment of Munc13. We develop β4-MC, derived from the Munc13-1 MUN and C2C domains, which bypasses Munc13 recruitment and directly mediates vesicle docking, priming and fusion after active zone disruption. In β4-MC, MUN domain functions are dispensable for docking, but active MUN and C2C domains are both necessary for neurotransmitter release. In wild-type mouse and human neurons, β4-MC but not β4-Zn strengthens neurotransmitter release. Furthermore, β4-Zn and β4-MC restore and enhance dopamine release in the mouse striatum, respectively. Hence, β4 proteins are engineered secretory proteins with therapeutic potential as enhancers of synaptic function.

## Introduction

Neurotransmitter release from presynaptic nerve terminals is essential for synaptic function. Activity-dependent changes in synaptic efficacy rely on plasticity of the release machinery and support information processing in the nervous system. Conversely, synaptic dysfunction is common in neurological disorders, and genes encoding presynaptic release machinery have been strongly implicated in disease ^1–25^. However, disease mechanisms are seldom monogenic, and multiple macromolecular machines regulate neurotransmitter release. Their mechanistic complexity and the large size of their proteins have impeded the development of new treatments, including gene therapy. For these reasons, few approaches exist to leverage neurotransmitter release as a control point for synaptic efficacy and as a therapeutic target to correct synaptic dysfunction.

Presynaptic active zones are specialized protein complexes that orchestrate neurotransmitter release. Multiple evolutionarily conserved protein families comprise the active zone and include RIM, RIM-BP, Liprin-α, ELKS, Munc13 and Piccolo/Bassoon ^26,27^. These proteins form multiple machineries to control secretion ^28–31^, and their coordinated activity primes and docks synaptic vesicles for exocytosis and mediates their coupling to voltage-gated Ca^2+^ channels (Ca_V_s) for rapid fusion during an action potential ^30–35^.

We previously engineered β4-Zn, a protein that bypasses the molecular complexity of the presynaptic active zone to mediate neurotransmitter exocytosis ^36^. β4-Zn is an 80 kDa protein composed of the RIM1 zinc finger and Ca_V_β4, a Ca_V_ subunit that confers targeting to the presynaptic plasma membrane. Simultaneous ablation of RIM and ELKS in mouse hippocampal neurons induces active zone disassembly ^31,37^, and in these neurons, β4-Zn reconstitutes synaptic vesicle priming, docking and action potential-evoked exocytosis without reassembling the megadalton-sized molecular machines that constitute the active zone ^36^. Mechanistically, the zinc finger of β4-Zn recruits the vesicle priming protein Munc13 to the presynaptic plasma membrane (Fig. S1A+B), and ablating this interaction disrupts β4-Zn activity ^33,36,38,39^. These findings inform a working model where β4-Zn recruits Munc13 to dock, prime and fuse vesicles with exceptional efficiency via the formation of exocytic SNARE complexes adjacent to Ca_V_s.

Here, we seek to design β4 secretory proteins that directly utilize Munc13 functional domains, bypassing Munc13 recruitment by β4-Zn. The main Munc13 proteins contain an N-terminal C2A domain followed by a linker and a calmodulin-binding motif, and C-terminal C1, C2B, MUN and C2C domains (Fig. S1A). Munc13 controls synaptic vesicle priming and fusion through SNARE complex assembly by its MUN domain ^34,35,40–46^. The C2A domain binds to RIM for active zone recruitment and Munc13 activation ^33,38,47,48^. Lipid binding C1, C2B and C2C domains flank the MUN domain, leading to a model in which the lipid binding domains connect the vesicular and plasma membranes and position the nested MUN domain for interactions with SNARE proteins ^49–54^ (Fig. S1B).

We report the development of an engineered secretory protein, β4-MC, from Ca_V_β4 and the Munc13-1 MUN and C2C domains, and we establish β4 proteins as enhancers of neurotransmitter release. In release-incompetent neurons with disrupted active zone assembly, β4-MC recovers synaptic vesicle priming, docking and exocytosis to reverse deficits in synaptic function. Perturbations of MUN or C2C domain function abolish β4-MC activity in release, but MUN domain activity in β4-MC is dispensable for vesicle docking. When expressed in wild-type mouse and human neurons, β4-MC boosts synaptic function. In the mouse dopaminergic system, β4-MC enhances striatal dopamine release, and β4-Zn restores release in dopamine neurons following RIM ablation. We conclude that β4-Zn and β4-MC are broadly applicable proteins for the recovery and enhancement of secretory function.

## Results

### β4-MC, an engineered protein of Ca_V_β4 and Munc13-1 MUN-C2C, mediates active zone function

We previously reconstituted neurotransmitter release in RIM and ELKS knockout neurons with β4-Zn (Figs. 1A, S1A+B). β4-Zn recruits endogenous Munc13 to the presynaptic plasma membrane to recover synaptic vesicle docking, priming and release ^36^. To design a protein that operates without a recruitment step, we directly incorporated essential Munc13 domains. We fused Ca_V_β4 to a Munc13-1 fragment containing the MUN and C2C domains (Figs. 1A, S1C-E), which regulate SNARE complex assembly ^40,42,46^ and bind to phospholipid membranes ^52,53,55^, respectively, thereby generating β4-MC.

**Figure 1.**
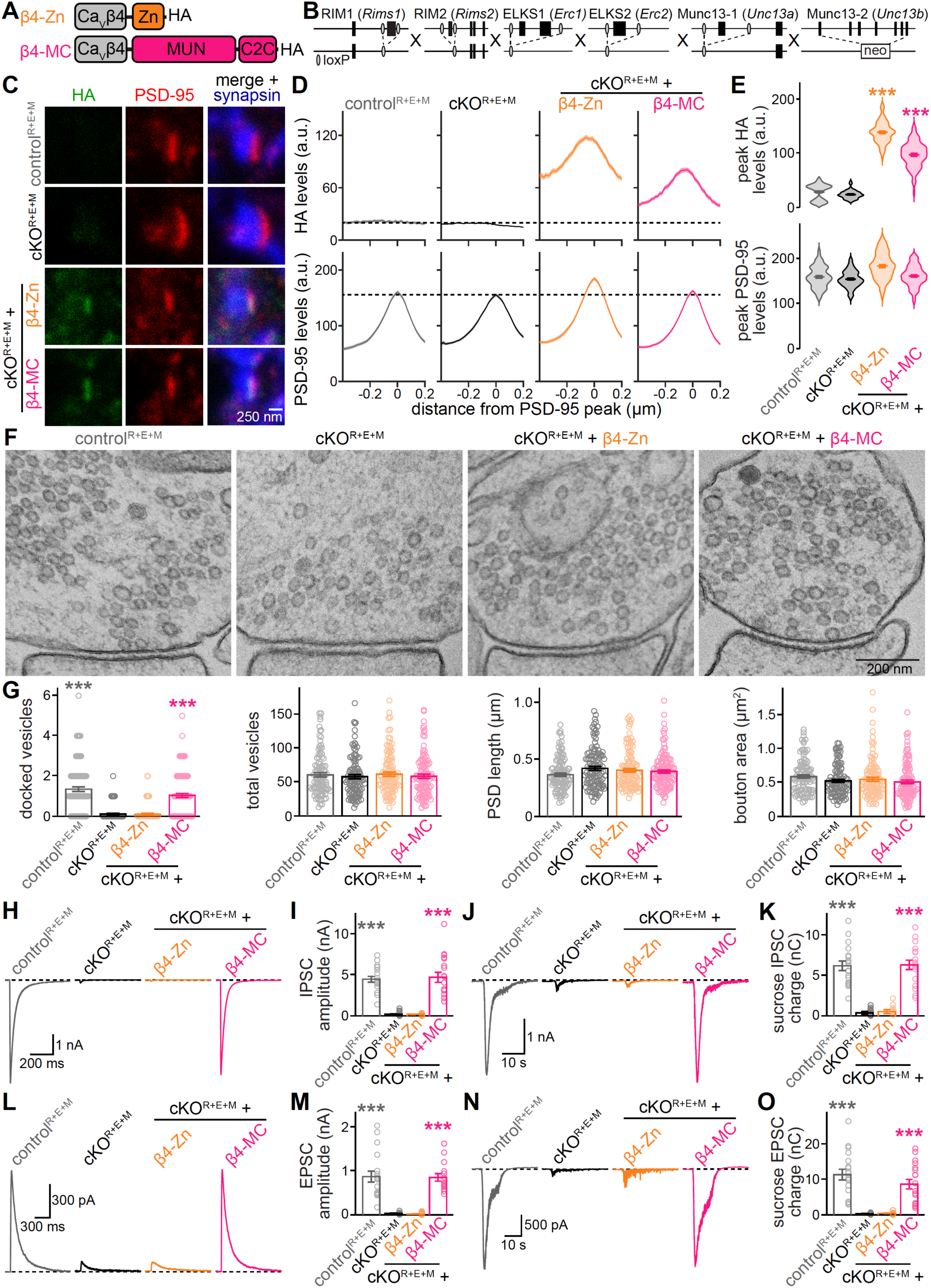
An engineered β4-MC protein of Ca_V_β4 and Munc13-1 MUN-C2C restores neurotransmitter release in RIM, ELKS and Munc13 knockout neurons. (A) Schematics of β4-Zn and β4-MC. (B) Strategy for simultaneous knockout of RIM1, RIM2, ELKS1, ELKS2, Munc13-1 and Munc13-_2_ 56. (C-E) Representative images (C) and summary plots of intensity profiles (D) and peak levels (E) of antibody signals at excitatory side-view synapses. Neurons were stained with antibodies against HA (imaged in STED), PSD-95 (STED), and synapsin (confocal). Dashed lines in D denote HA or PSD-95 levels in cKO^R+E+M^ neurons; control^R+E+M^ 84 synapses from 3 cultures, cKO^R+E+M^ 73/3, cKO^R+E+M^ + β4-Zn 80/3, cKO^R+E+M^ + β4-MC 63/3. (F and G) Representative electron micrographs of synapse cross-sections from high-pressure frozen neurons (F) and quantifications of docked vesicles, total vesicles, postsynaptic density (PSD) length, and bouton area (G); for docked vesicles and PSD length, control^R+E+M^ 110 active zones from 2 cultures, cKO^R+E+M^ 115/2, cKO^R+E+M^ + β4-Zn 118/2, cKO^R+E+M^ + β4-MC 116/2; for total vesicles and bouton area, control^R+E+M^ 100 synapses from 2 cultures, cKO^R+E+M^ 108/2, cKO^R+E+M^ + β4-Zn 105/2, cKO^R+E+M^ + β4-MC 105/2. (H and I) Representative traces (H) and quantification (I) of IPSCs recorded in whole-cell configuration and evoked by focal electrical stimulation; 18 cells from 3 cultures each. (J and K) As in H and I but for IPSCs evoked by local puffing of hypertonic sucrose; 18/3 each. (L and M) As in H and I but for electrically evoked NMDAR-mediated EPSCs; 18/3 each. (N and O) As in H and I but for sucrose-evoked AMPAR-mediated EPSCs; 18/3 each. Data are mean ± SEM; \*\*\**p* < 0.001, statistical significance compared to cKO^R+E+M^ was determined with Kruskal-Wallis tests followed by Dunn’s multiple comparisons post hoc tests in E, G, I, K, M, and O. For Munc13-1 sequences, additional expression analyses of engineered β4 proteins and Ca_V_2.1 localization in excitatory synapses, see Figure S1.

We then compared β4-Zn and β4-MC activity in a genetic model of active zone disassembly that also lacks endogenous Munc13 (Fig. 1B) ^56^. Specifically, cultured hippocampal neurons from mice with homozygous floxed alleles for the genes encoding RIM1, RIM2, ELKS1, ELKS2, and Munc13-1, and constitutive knockout alleles for the gene encoding Munc13-2 (Fig. 1B), were transduced with lentiviruses either expressing Cre recombinase to generate cKO^R+E+M^ neurons or expressing a truncated, inactive version of Cre to generate control^R+E+M^ neurons. β4-Zn and β4-MC were expressed in cKO^R+E+M^ neurons also by lentivirus, and they were hemagglutinin (HA)-tagged to facilitate detection. We have established before that active zone function is disrupted in these neurons following genetic ablation of RIM, ELKS and Munc13, and RIM and ELKS knockout leads to additional loss of RIM-BP and Piccolo/Bassoon ^31,36,56^.

To assess whether these engineered β4 proteins were targeted to the presynaptic plasma membrane in cKO^R+E+M^ neurons, we evaluated β4-Zn and β4-MC localization using STED microscopy. We selected side-view synapses that were identified by the presence of a synaptic vesicle cloud (stained with antibodies against synapsin, imaged in confocal) and an elongated postsynaptic density (stained with antibodies against PSD-95, STED) along the edge of the vesicle cloud following a workflow we established before ^36,57–59^. We then assessed β4-Zn and β4-MC localization (stained with antibodies against HA, STED) in line profiles that were drawn perpendicular to the postsynaptic density and quantified their peak levels (Fig. 1C-E). Both β4-Zn and β4-MC formed bar-like structures parallel to PSD-95 that peaked at ∼60 nm towards the presynaptic nerve terminal relative to PSD-95, establishing their localization at the presynaptic plasma membrane. There was a reduction in presynaptic Ca_V_2.1 in cKO^R+E+M^ neurons that was reversed by expression of either β4-Zn or β4-MC (Fig. S1F-H). We observed a similar Ca_V_2.1 phenotype before with β4-Zn in RIM and ELKS (cKO^R+E^) knockout neurons ^36^. These results indicate that β4-Zn and β4-MC are targeted to the presynaptic plasma membrane and promote presynaptic Ca^2+^ channel abundance in turn.

We next assessed whether β4-Zn and β4-MC recover synaptic vesicle docking, priming and release in cKO^R+E+M^ neurons with a combination of electron microscopy (Fig. 1F+G) and electrophysiology (Fig. 1H-O). For quantification of docking, we examined synapse ultrastructure in electron microscopic images of neurons following high-pressure freezing and freeze-substitution. Total synaptic vesicle counts, postsynaptic density length, and bouton area were unaffected in cKO^R+E+M^ neurons, but the number of docked synaptic vesicles (defined as those adjacent to the plasma membrane without visible space between the vesicular and plasma membranes) was strongly reduced ^56^. Strikingly, expression of β4-MC recovered vesicle docking to levels similar to those in control^R+E+M^ synapses. β4-Zn had no docking activity in cKO^R+E+M^ neurons (Fig. 1F+G), different from its effect in cKO^R+E^ neurons, in which β4-Zn fully restored docking ^36^.

We then recorded action potential-evoked inhibitory and excitatory postsynaptic currents (IPSCs and EPSCs) in response to electrical field stimulation (Fig. 1H+I, 1L+M) and estimated the pool of readily releasable vesicles (RRP) by application of hypertonic sucrose (Fig. 1J+K, 1N+O). Ablation of RIM, ELKS and Munc13 disrupted action potential- and sucrose-evoked PSCs at inhibitory and excitatory synapses, in agreement with previous findings ^56^. Compellingly, expression of β4-MC in cKO^R+E+M^ neurons restored IPSCs (Fig. 1H-K) and EPSCs (Fig. 1L-O), evoked either by action potentials or hypertonic sucrose. Release magnitude mediated by β4-MC was similar to that measured in control^R+E+M^ neurons. By contrast, β4-Zn did not restore any of these parameters, unlike its activity in cKO^R+E^ neurons ^36^.

Taken together, these results establish that β4-Zn is inactive in the absence of endogenous Munc13. In comparison, β4-MC performs essential active zone functions after ablation of RIM, ELKS and Munc13. Hence, β4-MC operates as a standalone protein to dock and prime synaptic vesicles for rapid, action potential-triggered neurotransmitter release.

### MUN-C2C is the minimally required Munc13-1 fragment for β4-MC activity

The Munc13 MUN domain is necessary for synaptic vesicle priming ^40–42,46^. However, it remains unclear whether the MUN domain alone can support neurotransmitter release, or whether flanking phospholipid-binding domains are needed to tether the MUN domain to the vesicular and plasma membranes ^41,49,52,54,60,61^. Taking these previous models into consideration, we devised two approaches to further refine β4-MC activity. First, we incorporated the entire C2B-MUN-C2C module into a β4 fusion protein, β4-CMC (Fig. 2A). Second, we generated β4 fusion proteins containing only C2C (β4-C, Fig. 2A) or only MUN (β4-M, Fig. S3A) to test whether they effectively enhance neurotransmitter release without their adjacent domain.

**Figure 2.**
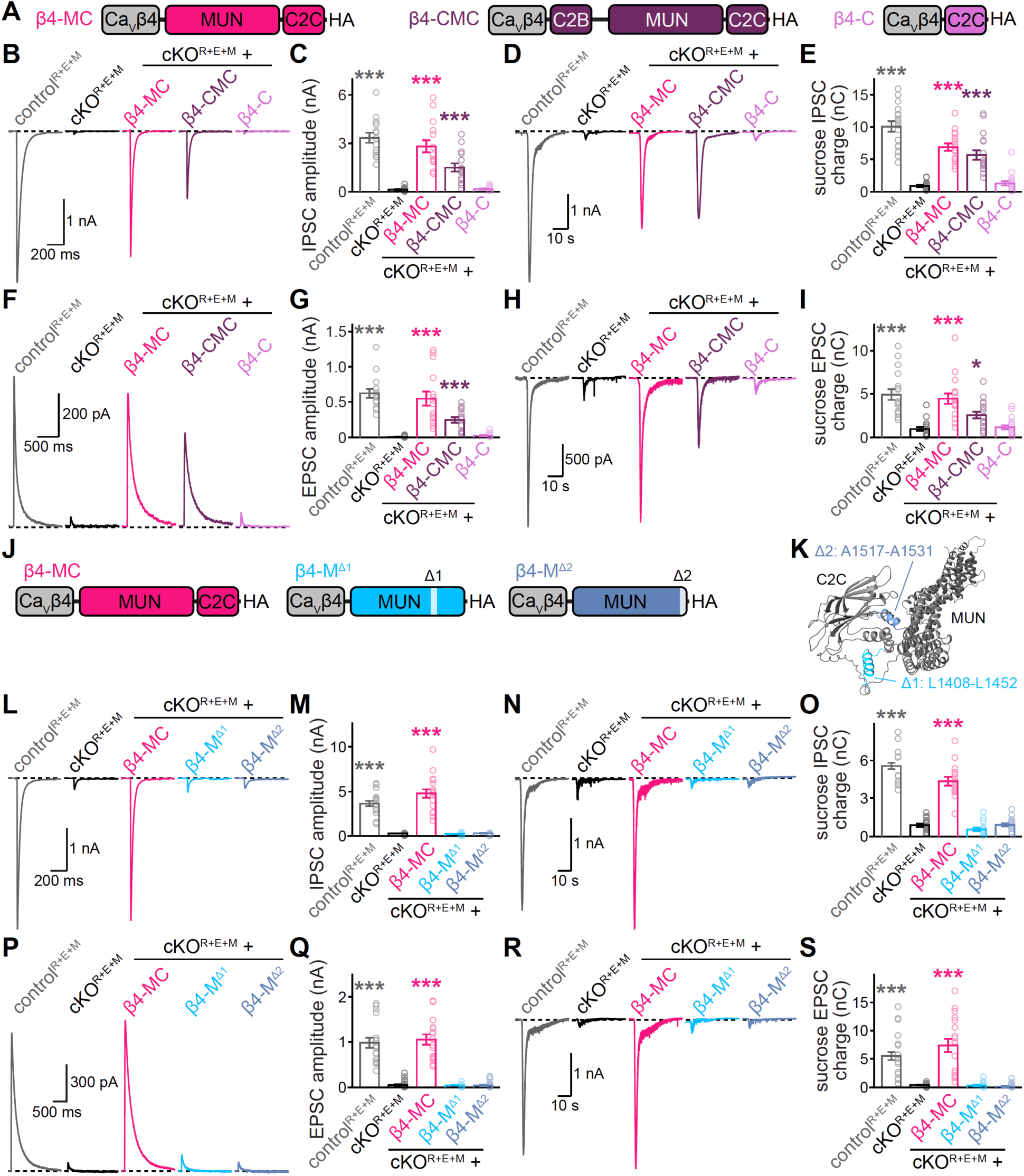
Removal of MUN or C2C from β4-MC abolishes rescue activity. (A) Schematics of β4-MC, β4-CMC and β4-C. (B and C) Representative traces (B) and quantification (C) of electrically evoked IPSCs from control^R+E+M^ neurons, cKO^R+E+M^ neurons, and cKO^R+E+M^ neurons following lentivirally mediated expression of the proteins in A; 15 cells from 3 cultures each. (D and E) As in B and C but for sucrose-evoked IPSCs; 17/3 each. (F and G) As in B and C but for electrically evoked NMDAR-mediated EPSCs; 15/3 each. (H and I) As in B and C but for sucrose-evoked AMPAR-mediated EPSCs; 17/3 each. (J) Schematics of β4-MC, β4-M^Δ1^ and β4-M^Δ2^. (K) MUN and C2C structure predicted by AlphaFold 3 indicating sequences removed for restoring protein expression. (L and M) Representative traces (L) and quantification (M) of electrically evoked IPSCs from control^R+E+M^ neurons, cKO^R+E+M^ neurons, and cKO^R+E+M^ neurons following lentivirally mediated expression of the proteins in J; 18 cells/3 cultures each. (N and O) As in L and M but for sucrose-evoked IPSCs; 18/3 each. (P and Q) As in L and M but for electrically evoked NMDAR-mediated EPSCs; 18/3 each. (R and S) As in L and M but for sucrose-evoked AMPAR-mediated EPSCs; 18/3 each. Data are mean ± SEM; \**p* < 0.05, \*\*\**p* < 0.001; statistical significance compared to cKO^R+E+M^ was determined with Kruskal-Wallis tests followed by Dunn’s multiple comparisons post hoc tests in C, E, G, I, M, O, Q and S. For additional expression and localization analyses of β4-MC, β4-CMC and β4-C, see Figure S2. For additional assessment of deletions within the MUN domain, see Figure S3.

Both β4-CMC and β4-C were efficiently expressed in transfected HEK293T cells and in lentivirally transduced neurons (Fig. S2A+B), and STED microscopy established their targeting to the presynaptic plasma membrane in cKO^R+E+M^ neurons (Fig. S2C-E). We next assessed whether β4-CMC and β4-C restored synaptic transmission in cKO^R+E+M^ neurons (Fig. 2B-I). β4-CMC recovered electrically and sucrose-evoked IPSCs (Fig. 2B-E) and EPSCs (Fig. 2F-I), though to a seemingly lesser extent than did β4-MC. Conversely, β4-C exhibited no rescue activity (Fig. 2B-I). In conclusion, addition of the C2B domain does not improve β4-MC efficacy and removal of the MUN domain abolishes it.

We next tested whether the MUN domain can operate without the neighboring C2C domain. We generated β4-M, but this construct failed to express in transfected HEK293T cells (Fig. S3A+B). Removal of a long loop (Δ1, residues L_1408_-L_1452_) or of a hydrophobic sequence (Δ2, residues A_1517_-A_1531_) might improve stability of the isolated MUN domain (Fig. 2J+K) ^42,51,62^. Deletions of both sequences as in β4-M^Δ1Δ2^ or β4-M^Δ1Δ2^C permitted expression in HEK293T cells and in lentivirally transduced neurons (Fig. S3C-E), but neither construct restored release in cKO^R+E+M^ neurons (Fig. S3F+G). We further found that the hydrophobic sequence, but not the long loop, was required for β4-MC activity by removing these sequences individually in β4-M^Δ1^C and β4-M^Δ2^C, though both constructs were expressed effectively (Fig. S3H-L). For a final comparison, we removed either the long loop (β4-M^Δ1^) or the hydrophobic stretch (β4-M^Δ2^) from β4-M. We then directly compared the expression (Fig. S3M+N), localization (Fig. S3O-Q) and activity (Fig. 2L-S) of these constructs to that of β4-MC. β4-M^Δ2^ was effectively localized to the presynaptic plasma membrane, but β4-M^Δ1^ had reduced presynaptic abundance (Fig. S3M-Q). However, neither β4-M^Δ1^ nor β4-M^Δ2^ restored synaptic responses evoked by action potentials or hypertonic sucrose (Fig. 2L-S). While weak expression and presynaptic localization might explain β4-M^Δ1^ inactivity, we overall conclude that MUN-C2C is the minimal sequence necessary for effective expression and restoration of neurotransmitter release.

Given that both MUN and C2C domains were important for β4-MC activity, we next tested whether their key molecular functions, SNARE complex assembly and phospholipid interactions, respectively, are needed (Fig. 3A+B). We produced β4-M^AA^C, in which we introduced mutations N_1128_A and F_1131_A in the MUN domain that eliminate its priming activity ^42^. We also inactivated the C2C domain using R_1598_E and F_1658_E mutations to disrupt lipid binding ^52,53^, generating β4-MC^EE^. To disrupt activities in both domains, we produced β4-M^AA^C^EE^. All constructs were effectively expressed in transfected HEK293T cells and transduced neurons (Fig. S4A+B), and they were targeted to the presynaptic plasma membrane (Fig. 3C-E).

**Figure 3.**
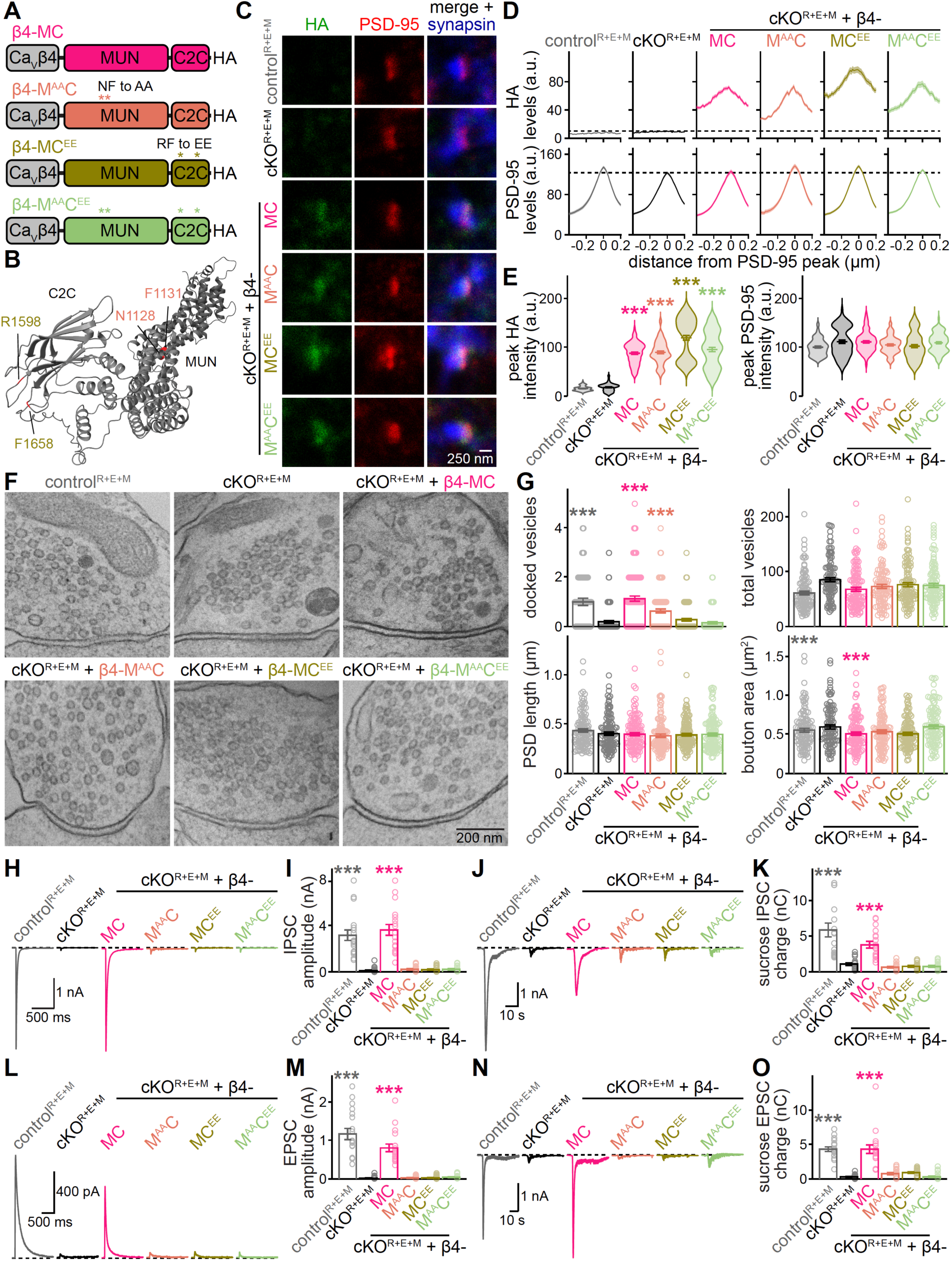
Inactivating mutations in MUN or C2C abolish β4-MC rescue activity. (A) Schematics of β4-MC, β4-M^AA^C, β4-MC^EE^ and β4-M^AA^C^EE^. (B) MUN and C2C structure predicted by AlphaFold 3 with residues mutated for inactivating domain functions indicated ^42,52,53^. (C-E) Representative images (C) and summary plots of intensity profiles (D) and peak levels (E) of antibody signals at excitatory side-view synapses stained for HA (STED), PSD-95 (STED) and synapsin (confocal). Dashed lines in D denote HA or PSD-95 levels in cKO^R+E+M^ neurons; control^R+E+M^ 81 synapses from 3 cultures, cKO^R+E+M^ 79/3, cKO^R+E+M^ + β4-MC 80/3, cKO^R+E+M^ + β4-M^AA^C 80/3, cKO^R+E+M^ + β4-MC^EE^ 78/3, cKO^R+E+M^ + β4-M^AA^C^EE^ 77/3. (F and G) Representative electron micrographs of synapse cross-sections from high-pressure frozen neurons (F) and quantifications (G); for docked vesicles and PSD length, control^R+E+M^ 109 active zones from 2 cultures, cKO^R+E+M^ 119/2, cKO^R+E+M^ + β4-MC 117/2, cKO^R+E+M^ + β4-M^AA^C 117/2, cKO^R+E+M^ + β4-MC^EE^ 113/2, cKO^R+E+M^ + β4-M^AA^C^EE^ 121/2; for total vesicles and bouton area, control^R+E+M^ 109 synapses from 2 cultures, cKO^R+E+M^ 118/2, cKO^R+E+M^ + β4-MC 117/2, cKO^R+E+M^ + β4-M^AA^C 116/2, cKO^R+E+M^ + β4-MC^EE^ 113/2, cKO^R+E+M^ + β4-M^AA^C^EE^ 120/2. (H and I) Representative traces (H) and quantification (I) of electrically evoked IPSCs; 18 cells from 3 cultures each. (J and K) As in H and I but for sucrose-evoked IPSCs; 18/3 each. (L and M) As in H and I but for electrically evoked NMDAR-mediated EPSCs; 18/3 each. (N and O) As in H and I but for sucrose-evoked AMPAR-mediated EPSCs; 18/3 each. Data are mean ± SEM; \*\*\**p* < 0.001; statistical significance compared to cKO^R+E+M^ was determined with Kruskal-Wallis tests followed by Dunn’s multiple comparisons post hoc tests in E, G, I, K, M, and O. For additional expression analyses of β4-MC, β4-M^AA^C, β4-MC^EE^ and β4-M^AA^C^EE^, see Figure S4.

Inactivating both the MUN and C2C domains as in β4-M^AA^C^EE^ ablated the docking (Fig. 3F+G) and release (Fig. 3H-O) activity of β4-MC in cKO^R+E+M^ neurons, consistent with essential roles for Munc13 in these processes ^34,44^. Remarkably, β4-M^AA^C, a protein with an inactive MUN domain, significantly enhanced vesicle docking in cKO^R+E+M^ neurons. However, these docked vesicles did not fuse in response to electrical stimulation or hypertonic sucrose (Fig. 3H-O), in line with a defect in synaptic vesicle priming following the loss of MUN domain function. β4-MC^EE^ did not restore docking (Fig. 3F+G) or vesicle fusion at inhibitory (Fig. 3H-K) or excitatory (Fig. 3L-O) synapses, indicating the need for C2C domain lipid binding in these activities of β4-MC.

These results establish that both the MUN and C2C domains are essential to β4-MC for neurotransmitter release. However, their functions in docking are separable: while C2C domain activity is needed for docking, MUN domain activity is dispensable. Hence, SNARE complex assembly might not be required for synaptic vesicle docking as assessed in high-pressure frozen neurons ^31,36,44,63^, at least not in the context of these β4 secretory proteins. Altogether, we conclude that β4-MC suffices to restore rapid, action potential-evoked neurotransmitter release and it requires functional MUN and C2C domains.

### β4-MC enhances neurotransmitter release from wild-type mouse and human neurons

We have thus far demonstrated that β4-Zn and β4-MC enhance neurotransmitter release in genetic models of active zone disassembly. Even so, simultaneous ablations of multiple active zone proteins as in these models are severe manipulations. To assess whether β4 secretory proteins enhance release in neurons with functional presynaptic release machinery, we characterized β4-Zn and β4-MC function in wild-type neurons.

Transduction of wild-type mouse hippocampal neurons with lentivirus expressing β4-Zn or β4-MC led to strong localization of both proteins at the active zone area of excitatory and inhibitory synapses (Fig. 4A-C, S4C-F). Strikingly, β4-MC increased vesicle docking and action potential-evoked transmission in wild-type neurons, though it did not alter sucrose-induced responses (Fig. 4D-I). In contrast, expression of β4-Zn had no detectable effect on docking or release (Fig. 4D-I), and neither construct changed the active zone abundance of Ca_V_2.1 (Fig. S4G-L). Rather, we found a decrease in the ratio of release evoked by paired pulses in neurons expressing β4-MC, indicating that β4-MC increased vesicular release probability (Fig. 4J+K).

**Figure 4.**
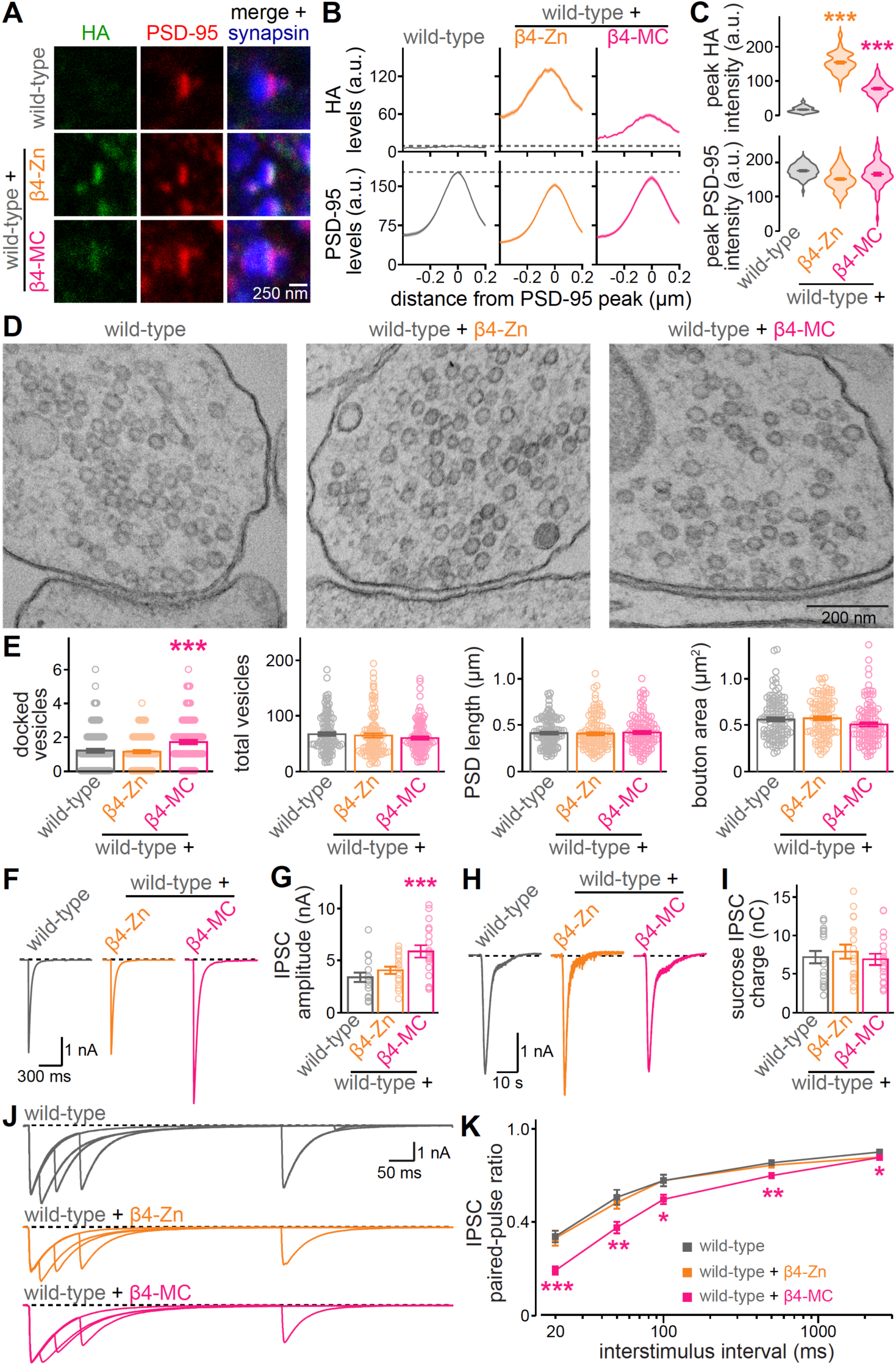
β4-MC enhances docking and neurotransmitter release. (A-C) Representative images (A) and summary plots of intensity profiles (B) and peak levels (C) of antibody signals at excitatory side-view synapses of hippocampal wild-type neurons and wild-type neurons transduced with β4-Zn or β4-MC. Neurons were stained for HA (STED), PSD-95 (STED) and synapsin (confocal). Dashed lines in B denote HA or PSD-95 levels in wild-type neurons; wild-type 74 synapses from 3 cultures, wild-type + β4-Zn 83/3, wild-type + β4-MC 70/3. (D and E) Representative electron micrographs of synapse cross-sections from high-pressure frozen neurons (D) and quantifications (E); for docked vesicles and PSD length, wild-type 113 active zones from 2 cultures, wild-type + β4-Zn 106/2, wild-type + β4-MC 104/2; for total vesicles and bouton area, wild-type 113 synapses from 2 cultures, wild-type + β4-Zn 106/2, wild-type + β4-MC 104/2. (F and G) Representative traces (F) and quantification (G) of electrically evoked IPSCs; 18 cells from 3 cultures each. (H and I) As in F and G but for sucrose-evoked IPSCs; 18/3 each. (J and K) As in F and G but for paired-pulse ratios of electrically evoked IPSCs; 18/3 each. Data are mean ± SEM; \**p* < 0.05, *\*\*p* < 0.01, \*\*\**p* < 0.001, statistical significance compared to wild-type was determined with Kruskal-Wallis tests followed by Dunn’s multiple comparisons post hoc tests in C, E, and G, and by a two-way ANOVA (genotype *p* < 0.001, interstimulus interval *p* < 0.001, interaction *p* = 0.001) followed by Dunnett’s multiple comparisons test (shown in figure) in K. For additional expression and localization analyses of β4-Zn and β4-MC, and for Ca_V_2.1 localization at wild-type excitatory and inhibitory synapses, see Figure S4.

β4 proteins might have therapeutic potential as enhancers of neurotransmitter release. To determine whether they act as release enhancers in human neurons, we derived cortical glutamatergic neurons from KOLF2.1J human induced-pluripotent stem cells (hiPSCs) by expression of Neurogenin-2 (Fig. 5A) ^63^. Lentiviral transduction resulted in effective expression of β4-Zn and β4-MC in human neurons (Fig. 5B). As in cultured mouse hippocampal neurons, β4-MC but not β4-Zn increased release induced by electrical stimulation, while the pool of vesicles released in response to hypertonic sucrose was not measurably changed (Fig. 5C-F). In summary, β4-MC enhances release at wild-type synapses in mouse and human neurons.

**Figure 5.**
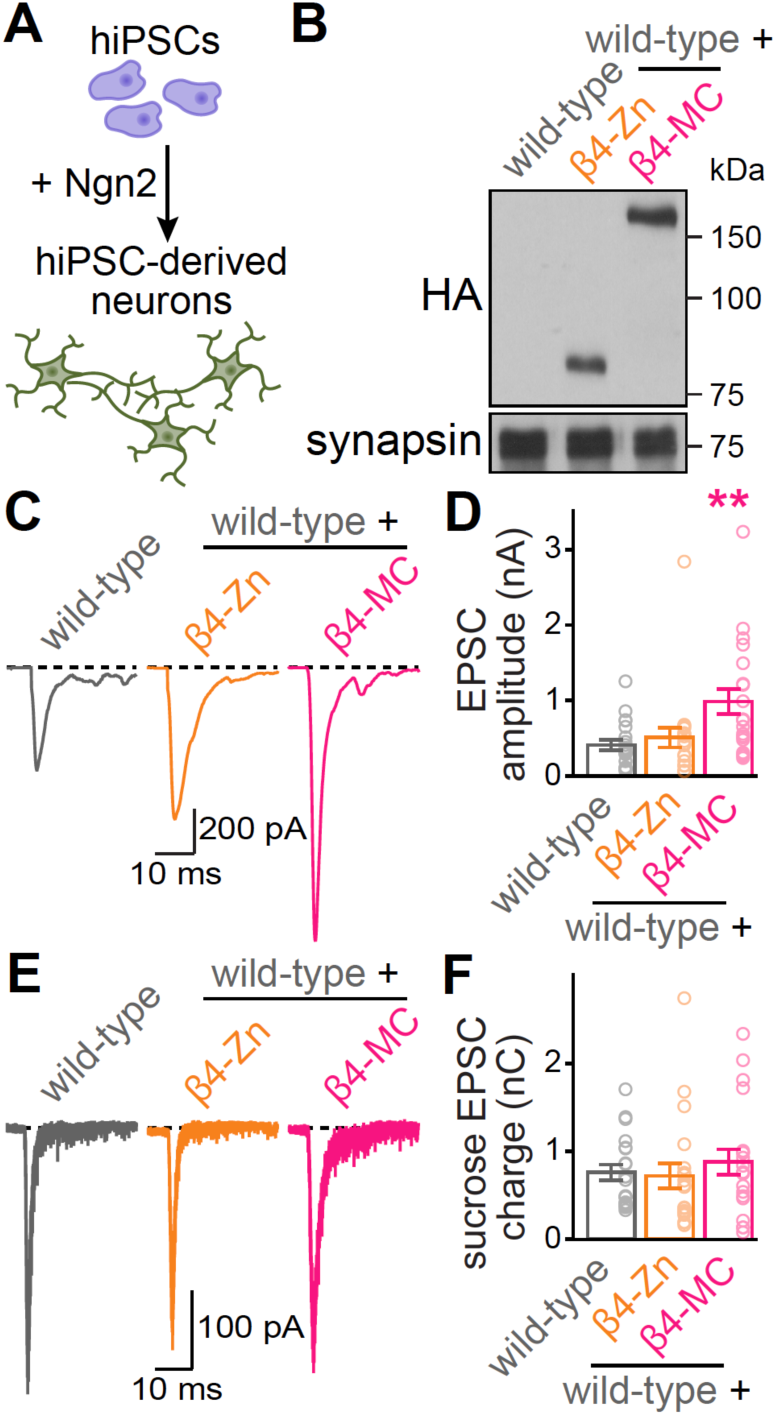
β4-MC enhances neurotransmitter release in human neurons. (A) Experimental workflow; human induced pluripotent stem cells (hiPSCs) were differentiated into excitatory neurons with expression of Neurogenin-2 ^63^. (B) Western blot for HA and synapsin from homogenates of cultured hiPSC-derived neurons and of hiPSC-derived neurons following lentivirally mediated expression of β4-Zn or β4-MC; a representative blot from two repeats is shown. (C and D) Representative traces (C) and quantification (D) of electrically evoked AMPAR-mediated EPSCs; 20 cells from 2 differentiations each. (E and F) As in C and D but for sucrose-evoked AMPAR-mediated EPSCs; 20/2 each. Data are mean ± SEM; \*\**p* < 0.01; statistical significance compared to wild-type was determined with Kruskal-Wallis tests followed by Dunn’s multiple comparisons post hoc tests in D.

### Designing β4 fusion proteins for conditional, AAV-based in vivo use

Adeno-associated viruses (AAVs) are standard vectors for delivery of exogenous genetic material in vivo. AAVs are limited in their genetic load to approximately 5 kilobases, which is exceeded by the length of the β4-MC expression cassette. As β4-MC contains the smallest Munc13 fragment for mediating neurotransmitter release, we optimized the sizes of Ca_V_β4 and of the promoter driving expression.

Ca_V_β auxiliary subunits contain an SH3 domain, a GK domain, and additional N-terminal and C-terminal segments, with the GK domain forming the binding pocket for pore-forming Ca_V_α subunits ^64^. To identify the minimal Ca_V_β4 sequences necessary for active zone targeting, we generated two shorter variants of Ca_V_β4 and fused them to the RIM1 zinc finger domain (Fig. S5A). In β4^GK^-Zn, we removed all Ca_V_β4 sequences except for the GK domain. In β4^BID-GK^-Zn, we removed all Ca_V_β4 sequences except for the GK domain and a short preceding sequence known as the beta-interacting domain (BID) that may stabilize the GK domain. β4^GK^-Zn and β4^BID-GK^-Zn were expressed in transfected HEK293T cells and transduced neurons, but initial assessment of their function suggested that neither mediated neurotransmitter release in cKO^R+E^ neurons (Fig. S5B-E). We next took a more moderate approach and designed Ca_V_β4 variants based on Ca_V_β splice isoforms (Fig. S5F), removing some of the Ca_V_β4 N-terminus to generate β4^M^-MC (“β4^M^” for medium) or most of the N- and C-termini to generate β4^S^-MC (“β4^S^” for short). β4^M^-MC and β4^S^-MC were expressed similarly in transfected HEK293T cells and transduced neurons (Fig. S5G+H), and both appeared to recapitulate the release enhancing phenotype of β4-MC in wild-type mouse hippocampal neurons (Fig. S5I+J). Hence, we proceeded with β4^S^.

We cloned cDNAs for β4^S^-Zn and β4^S^-MC into a double floxed inverted (DIO) expression cassette, with expression driven by a synthetic, short promoter (SP) ^65^ to generate SP^DIO^-β4^S^-Zn and SP^DIO^-β4^S^-MC. For comparison of promoter efficacy, we also generated hSyn^DIO^-β4^S^-Zn, for which expression is driven by the human synapsin (hSyn) promoter used in all other lentiviral vectors thus far (Fig. S6A). Expression efficacy of SP in transfected HEK293T cells and in lentivirally transduced cKO^R+E^ neurons was comparable to that of hSyn (Fig. S6B+C). Moreover, hSyn^DIO^-β4^S^-Zn, SP^DIO^-β4^S^-Zn and SP^DIO^-β4^S^-MC restored neurotransmitter release evoked by action potentials in cKO^R+E^ neurons (Fig. S6D+E). With these modifications, we established AAV-compatible, conditional expression cassettes.

### Engineered β4 proteins enhance striatal dopamine release

Neuromodulatory systems are often affected in neurological disease. A hallmark example is dopamine neurons, many but not all of which degenerate in Parkinson’s Disease (PD) ^66–70^. Hence, we tested whether β4 proteins can be used to enhance dopamine release, first in mice with genetic ablation of RIM in dopamine neurons (RIM cKO^DA^ mice, Fig. 6A). Genome-wide sequencing has identified RIM proteins as risk and disease progression factors in PD ^9,10^, and we have previously established that RIM mediates dopamine release ^71–74^. We used cre-dependent expression of β4-Zn selectively in dopamine neurons in RIM cKO^DA^ mice (Fig. 6B). Unilateral stereotaxic delivery of hSyn^DIO^-β4^S^-Zn into the substantia nigra (SNc) led to strong expression of β4^S^-Zn in dopamine axons in the ipsilateral striatum (Fig. S7A-C). We then monitored striatal dopamine release in acute brain slices prepared from mice that also expressed the fluorescent dopamine sensor GRAB_DA3m_ in the dorsal striatum ^73,75,76^. Dopamine axons were stimulated with a focal stimulation electrode, and we used single stimuli, 10 Hz 10 stimulus trains, and 1 Hz stimulation for 130 s to evoke dopamine release. RIM ablation impaired dopamine release when assessed with GRAB_DA3m_, as we have established before with other sensors and methodology ^71–74^. Expression of β4^S^-Zn greatly enhanced striatal dopamine release in RIM cKO^DA^ mice to levels nearing those measured in RIM control mice. This release enhancement was observed for single stimuli and stimulus trains (Fig. 6C-E) and was also maintained during prolonged stimulation at 1 Hz (Fig. S7D+E). Hence, β4^S^-Zn reverses release deficits in dopamine neurons that lack RIM.

**Figure 6.**
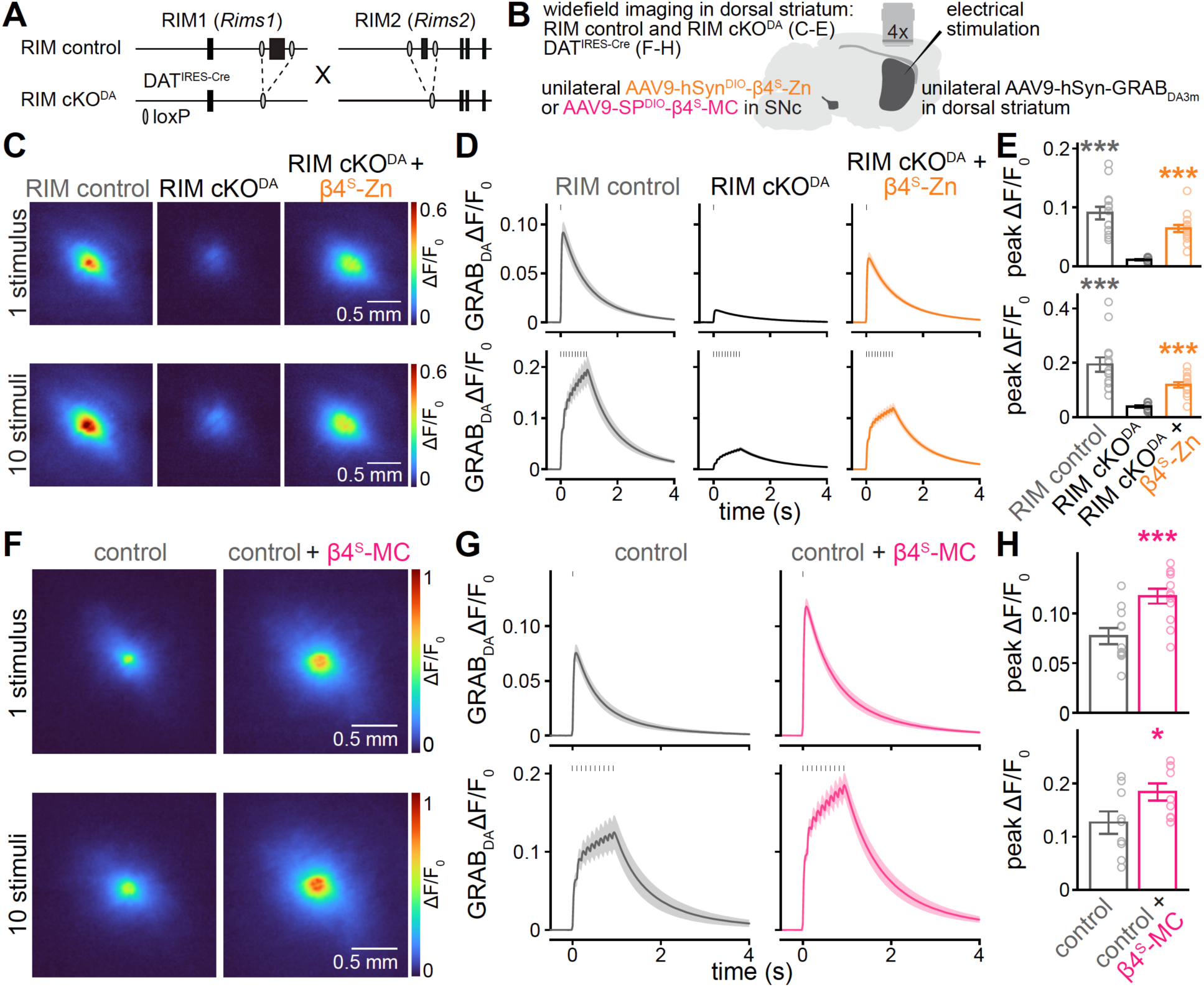
β4-Zn and β4-MC enhance release from dopamine neurons. (A) Strategy for knockout of RIM1 and RIM2 in dopamine neurons ^71^. (B) Experimental workflow; AAV9-hSyn^DIO^-β4^S^-Zn was injected unilaterally into the right substantia nigra (SNc) of RIM cKO^DA^ mice, or AAV9-SP^DIO^-β4^S^-MC was injected unilaterally into the right SNc of DAT^IRES-Cre^ (control) mice. Striatal dopamine release in the same hemisphere was monitored via GRAB_DA3m_ imaging in acute sagittal slices in the presence of 1 µM DHβE four to five weeks after injection. (C-E) Representative images of peak GRAB_DA3m_ fluorescence (C) evoked by a single electrical stimulus or by 10 stimuli at 10 Hz in the dorsolateral striatum of RIM control mice, RIM cKO^DA^ mice or RIM cKO^DA^ mice expressing β4^S^-Zn, and quantification of the GRAB_DA3m_ ΔF/F_0_ time course (D) and peak ΔF/F_0_ (E). Vertical ticks in E denote application of electrical stimulation; for 1 stimulus, RIM control 14 slices from 4 mice, RIM cKO^DA^ 13/4, RIM cKO^DA^ + β4^S^-Zn 15/4; for 10 stimuli at 10 Hz, RIM control 14/4, RIM cKO^DA^ 13/4, RIM cKO^DA^ + β4^S^-Zn 14/4. (F-H) As in C-E but for GRAB_DA3m_ fluorescence evoked by a single stimulus or by 10 stimuli at 10 Hz in control mice or control mice expressing β4^S^-MC, all mice had a DAT^IRES-Cre^ allele; for 1 stimulus, 12/4 each; for 10 stimuli at 10 Hz, 9/3 each. Data are mean ± SEM. \**p* < 0.05, \*\*\**p* < 0.001, statistical significance compared to RIM cKO^DA^ in E was determined with Kruskal-Wallis tests followed by Dunn’s multiple comparisons post hoc tests. Statistical significance compared to control in H was determined with Mann-Whitney tests. For the generation of short Ca_V_β4 sequences, see Figure S5. For the generation of double floxed inverted (DIO) expression vectors for β4 proteins, see Figure S6. For quantification of β4 protein expression in the dorsolateral striatum and of dopamine release evoked by 1 Hz stimulus trains, see Figure S7.

Enhancing dopamine signaling with L-DOPA, a dopamine precursor, is the gold standard therapy for PD ^66,77^. While effective in improving bradykinesia, prolonged L-DOPA administration leads to side effects, and other symptoms associated with PD including cognitive decline may be less amenable to L-DOPA treatment. One strategy to improve PD treatment might be to enhance dopamine release from surviving dopamine neurons, which might restore striatal dopamine levels and dopamine dynamics. Hence, we tested whether engineered β4 proteins operate as release enhancers in wild-type dopamine neurons. We specifically evaluated β4^S^-MC, given its release enhancement phenotype in cultured mouse and human neurons (Figs. 4, 5, S5). We injected AAV9 expressing SP^DIO^-β4^S^-MC unilaterally into the SNc of DAT^IRES-Cre^ mice and found specific expression in ipsilateral striatal dopamine axons (Fig. S7F-H). Remarkably, expression of β4^S^-MC increased GRAB_DA3m_ transients in response to single and train stimuli (Fig. 6F-H, Fig. S7I+J). Hence, β4-MC enhances dopamine release in the mouse striatal dopamine system.

Overall, our work establishes that β4-Zn and β4-MC can be used to restore and enhance release across a variety of preparations that include glutamate, GABA and dopamine neurons, cultured neurons and intact striatal circuits, and mouse and human neurons.

## Discussion

Engineered β4 proteins are compact secretory proteins that restore and strengthen neurotransmission. We described the RIM-based fusion enhancer β4-Zn before ^36^, and we here develop and establish Munc13-derived β4-MC. β4-MC contains the Munc13-1 MUN and C2C domains and mediates vesicle docking, priming and release. While β4-MC with perturbed MUN domain activity can dock synaptic vesicles, it requires active MUN and C2C domains for neurotransmitter release. We then assess the activities of β4-Zn and β4-MC in multiple model systems and optimize them for in vivo use. Notably, β4-Zn and β4-MC recover synaptic transmission in neurons with disrupted active zones, and β4-MC also enhances release in wild-type neurons. We demonstrate β4-Zn and β4-MC function in excitatory and inhibitory mouse hippocampal neurons, in glutamatergic human neurons, and in the mouse striatal dopamine system. Our work establishes Munc13 MUN and C2C domains as a minimal fusion-promoting mechanism and develops their release-enhancing function towards therapeutic use.

### Minimal Munc13 domains for synaptic vesicle priming and docking

The Munc13 MUN domain is necessary for synaptic vesicle priming and fusion. It initiates the opening of Syntaxin, which enables SNARE complex assembly of Syntaxin, SNAP-25 and VAMP/synaptobrevin ^40–42,45,46,78^. The exact contributions of other Munc13 domains remain to be defined. One model suggests that the Munc13 C-terminal C1-C2B-MUN-C2C domains form a membrane bridge between synaptic vesicles and the presynaptic membrane ^52,55,79^, positioning the MUN domain for interactions with SNARE proteins. Activity-dependent changes in C1 or C2B phospholipid binding may reorient the MUN domain for progression through SNARE complex assembly ^54,60^. However, in adrenal chromaffin cells and at the C. elegans neuromuscular junction, Munc13 proteins without C1 or C2B efficiently prime vesicles ^53,80,81^, suggesting that these domains are not essential. Evidence from in vitro fusion assays indicates that the C2C domain may be similarly dispensable ^45,55^. Indeed, Munc13 fragments containing only the MUN domain may be sufficient for priming ^41,45^. Here, we characterize the functional activity of Munc13 domains in a simplified molecular context through expression of engineered β4 proteins following active zone disassembly, and we show that synaptic vesicle priming requires at least functional MUN and C2C domains, as in β4-MC (Figs. 1-3). Notably, β4-MC does not include the C2B domain, and inclusion of the C2B domain (in β4-CMC, Fig. 2) did not measurably enhance priming. For β4 proteins, direct tethering to Ca^2+^ channel α subunits may serve as a membrane anchor, rendering the anchoring mechanisms of C1 and C2B redundant. In addition, the C1 and C2B domains can be autoinhibitory ^82^, and their absence may contribute to the apparent increase in efficacy of β4-MC relative to β4-CMC.

The relationship between synaptic vesicle docking and priming has been debated ^31,36,44,83–85^. Docking and priming might represent equivalent states, and the same molecular mechanism might generate fusion-competent synaptic vesicles and anchor vesicles to the presynaptic plasma membrane. Alternatively, docking and priming might rely on distinct mechanisms, and addition of a vesicle to the RRP may not always correlate with docking to the presynaptic plasma membrane. Here, we found that β4-MC docks synaptic vesicles even after inactivation of the MUN domain (Fig. 3). Hence, Munc13 C2C domain activity in β4-MC suffices to dock synaptic vesicles, and MUN-mediated SNARE complex assembly might be dispensable for the tight attachment of these vesicles to the presynaptic plasma membrane. An alternative explanation might be that the MUN domain mutations are too weak to fully disrupt docking, though they are sufficient to ablate the release function of β4-MC. We conclude that synaptic vesicles can be docked but not primed, at least after disrupting active zone assembly. Conversely, we have previously found that genetic disassembly of the active zone abolishes docking, but some synaptic vesicle priming persists ^31^. Together, these findings support the model that vesicle docking and priming can be mediated by separable molecular processes.

### Mechanistic comparison of β4-Zn and β4-MC

β4-Zn and β4-MC differentially modulate synaptic function. In wild-type neurons, β4-MC enhances synaptic vesicle docking and neurotransmitter release, but β4-Zn has no effect on synaptic efficacy (Fig. 4). Because β4-Zn operates through endogenous Munc13 (Fig. 1) ^36^ whereas β4-MC directly utilizes Munc13 functional domains, several possible explanations might account for the difference in enhancement. First, Munc13 clusters are the molecular correlates of presynaptic release sites where synaptic vesicles dock, prime and fuse ^54,86,87^. β4-MC might promote the formation of new release sites near Ca^2+^ channels. This mechanism might be particularly powerful in dopamine axons in which many varicosities might not contain a full set of release site proteins ^71,72,88^. Conversely, β4-Zn might compete with endogenous RIM for recruitment of Munc13, and Munc13 abundance is likely rate-limiting for β4-Zn efficacy. Second, absence of autoinhibitory C1 and C2B domains in β4-MC might accelerate SNARE complex assembly and promote vesicle fusion ^54,60,82^. Third, with β4-MC, a priming module is directly tethered to Ca_V_s, which might reduce coupling distance between fusion-competent synaptic vesicles and Ca^2+^ sources, enhancing vesicular release probability. These data are consistent with studies in C. elegans, where expression of similar Munc13 fragments also enhances release probability ^82,89^.

β4-MC increases striatal dopamine release evoked by short stimulus trains (Fig. 6), suggesting that β4-MC upregulates the availability of releasable vesicles and release sites. The midbrain dopaminergic system forms dense axonal arborizations in the striatum. Recent work indicates that only a fraction of the axonal varicosities contains active zone-like release sites and that these release sites are responsible for most striatal dopamine release ^71–73,88,90^. In contrast, most axonal varicosities have action potential-induced Ca^2+^ entry ^88^. Taking these findings into consideration along with the seemingly broad localization of β4-Zn and β4-MC in striatal dopamine axons (Fig. S7), Ca_V_s may be widespread in these axons. β4-MC might form new release sites in dopamine axons by recruiting and priming dopamine-containing vesicles adjacent to axonal Ca_V_s.

### Therapeutic potential of β4 fusion proteins

Dysfunctions of active zone proteins, other proteins in the presynaptic release machinery, and secretion in general, have been increasingly associated with neurological disease ^1–25^. β4 proteins may have therapeutic potential to correct secretory deficits or to modulate circuit function, for example for disorders that involve active zone protein dysfunction. *UNC13A*, the gene encoding Munc13-1, is a major genetic risk factor for TDP-43-associated forms of amyotrophic lateral sclerosis and frontotemporal dementia ^1–4^, which may account for ∼97% and ∼45% of sporadic cases, respectively. TDP-43 pathology induces mis-splicing of *UNC13A* mRNA, leading to depletion of presynaptic Munc13 and impairment of synaptic function ^3–5,91^. Munc13 loss of function has also been implicated in spinal muscular atrophy and neurodevelopmental syndromes ^92,93^. Because Munc13 does not fit within an AAV for gene therapy, β4-MC or proteins similar to it could be used to restore Munc13 functions in these disorders.

Another example is PD, in which dopamine neurons in the substantia nigra pars compacta degenerate. *RIMS1* and *RIMS2*, the genes encoding human RIM proteins, were identified as risk and disease progression factors in PD ^9,10^, respectively. We demonstrate that β4-MC enhances dopamine release, and we speculate that β4-MC expression might be used to counteract dopamine loss in PD by enhancing release from surviving dopamine neurons and restoring dopamine dynamics. Because L-DOPA treatment broadly increases striatal dopamine levels, prolonged treatment induces dyskinesia and other side effects in most patients ^66,94^. Enhancement of dopamine release through β4-MC may present a viable alternative. Future efforts should evaluate β4-Zn and β4-MC efficacy as release enhancers across models of PD.

A key advance in our work is the optimization of β4-Zn and β4-MC for expression via AAVs, an essential step towards gene therapy. Because β4 proteins act cell-autonomously, combination of their release enhancing function with targeted delivery, for example by using cell-type specific enhancers ^95–98^, may be a general strategy to restore nervous system function. These strategies could be applied to enhance excitation via glutamate neurons, to enhance inhibition via GABA neurons, or to tune circuit function by enhancing neuromodulatory transmitter signaling, as we establish for dopamine. Finally, β4 proteins might be broadly active in other systems with regulated exocytosis. Examples include transmitter release in the peripheral sensory systems, the secretion of monoamines from the adrenal gland, and the secretion of hormones such as insulin. While the molecular requirements might vary by system, the functionality of β4 proteins across multiple neuron types, transmitter systems and species suggests broad applicability of the framework we present to engineer fusion-enhancing proteins.

## Acknowledgments

We thank members of the Kaeser laboratory for insightful discussions. We acknowledge F. Ureña Ortiz, E. Butler and G. Handy for technical support, and N. Brose and C. Imig for sharing Munc13-1 and Munc13-2 mutant alleles. We thank R. Held, A. Brunger and J. Rizo for discussions on the design of engineered β4 proteins; Z. Pang, S. D. de León-Guerrero and X. Li for advice on differentiating human neurons; G. de Nola for advice on electron microscopy; and, M. Grompe and S. Chai for advice on the use of the short promoter. This work was supported by the NIH (R01MH113349, R01NS083898, R01DA056109, R01NS103484 to P.S.K. and F31NS127399 to M.C.), Stuart H.Q. and Victoria Quan fellowships (to M.C. and R.C.L.), and Harvard Medical School. K.Y. was supported by a Human Frontier Science Program postdoctoral fellowship (LT0023/2025-L). R.C.L. is the recipient of a National Science Foundation graduate research fellowship (DGE2140743). C.T. is currently at Fudan University. We acknowledge the Boston Children’s Hospital Viral Core, the Neurobiology Imaging Facility (NIF), the Core for Imaging Technology and Education (CITE), and the Electron Microscopy Core at Harvard Medical School.

## Author contributions

Conceptualization: MC, KY, RCL, CT, PSK; Methodology: MC, KY, RCL, CT; Formal analysis: MC, KY, RCL, CT, PSK; Investigation: MC, KY, RCL, CT; Resources: MC, CT; Writing – original draft: MC, PSK; Writing – review and editing: MC, KY, RCL, CT, PSK; Supervision: PSK; Funding: PSK

## Methods

### Mice

Mice for simultaneous ablation of RIM1, RIM2, ELKS1, ELKS2, Munc13-1 and Munc13-2 were described previously ^56^ and contain homozygous floxed alleles for *Rims1* (to remove RIM1α and RIM1β), *Rims2* (to remove RIM2α, RIM2β and RIM2γ), *Erc1* (to remove ELKS1α), *Erc2* (to remove ELKS2α), *Unc13a* (to remove Munc13-1), and homozygous constitutive knockout alleles for *Unc13b* (to remove Munc13-2). For conditional quadruple knockout of RIM1, RIM2, ELKS1 and ELKS2, mice with homozygous floxed alleles for each were used and were described previously ^31,36^. For experiments in mice or neurons without gene ablation, mice with floxed alleles were used without the addition of Cre recombinase, or Cre mice without floxed alleles were used. For mice with expression of Cre recombinase in dopamine neurons, mice heterozygous for DAT^IRES-Cre^ were used ^99^. For conditional knockout of RIM1 and RIM2 in dopamine neurons, mice with floxed alleles for *Rims1* and *Rims2* were crossed to DAT^IRES-Cre^ mice as described before ^71,73,99^. RIM cKO^DA^ mice were homozygous for floxed RIM1 and RIM2 and heterozygous for DAT^IRES-Cre^, and RIM control mice were heterozygous for floxed RIM1 and RIM2 and heterozygous for DAT^IRES-Cre^. Mice were housed as breeding pairs or separated by sex in a mouse breeding facility maintained in rooms set to 22 °C (range: 20–24 °C) and 50% humidity (range: 35–70%) under a regular light–dark cycle, with free access to food and water. Animal experiments were performed according to approved protocols at Harvard University.

### HEK293T cells

HEK293T cells, an immortalized cell line, were cultured as described before ^28,36,57,59^. They were purchased from ATCC (CRL-3216, RRID: CVCL_0063), expanded, and stored in liquid nitrogen until use. After thawing, cells were maintained in Dulbecco’s Modified Eagle Medium supplemented with 10% fetal bovine serum (Atlas Biologicals F-0500-D) and 1% penicillin-streptomycin. HEK293T cells were passaged upon reaching 70-80% confluency every 1 to 3 d and replaced from the frozen stock after approximately 20 passages.

### Primary mouse hippocampal neurons

Primary mouse hippocampal cultures were generated from newborn mice as described before ^28,36,57,59^. Hippocampi were harvested from newborn mice of either sex within 24 h after birth and dissociated. Dissociated cells were plated onto 12 mm glass coverslips (for STED microscopy, electrophysiology or western blotting) or 6 mm sapphire discs (Technotrade International 616-100, for electron microscopy) coated with Matrigel. For ∼24 h after plating, cultures were maintained in plating medium composed of Minimum Essential Medium (MEM) supplemented with 0.5% glucose, 0.02% NaHCO_3_, 0.1 mg/mL transferrin, 10% Fetal Select bovine serum (Atlas Biologicals FS-0500-AD), 2 mM L-glutamine, and 25 μg/mL insulin. After ∼24 h, the plating medium was exchanged for growth medium composed of MEM supplemented with 0.5% glucose, 0.02% NaHCO_3_, 0.1 mg/mL transferrin, 5% Fetal Select bovine serum (Atlas Biologicals FS-0500-AD), 2% B-27 supplement (Thermo Fisher 17504044), and 0.5 mM L-glutamine. On day in vitro (DIV) 1 to 2, depending on growth, 50-75% of the growth medium was exchanged for growth medium supplemented with 2-4 μM cytosine arabinoside (AraC). Cultures were maintained in a 37 °C tissue culture incubator until DIV15-20.

### Human induced pluripotent stem cells

Human induced pluripotent stem cells (hiPSCs) from the KOLF2.1J line were purchased from JAX (JIPSC001000), expanded, and stored in liquid nitrogen until use following established protocols ^100,101^. After thawing, cells were plated onto Matrigel-treated 35 mm round bottom dishes and maintained in mTeSR Plus medium (STEMCELL 100-0276) supplemented with 100 µg/mL Primocin. Upon reaching 60-80% confluency, hiPSCs were detached with Accutase (STEMCELL 07922) and passaged for a maximum of 15 passages. For ∼24 h after thawing and after each passage, hiPSC culture medium was supplemented with CEPT cocktail comprising: 50 nM Chroman, 5 µM Emricasan, 0.7 µM trans-ISRIB and 1 µL/mL polyamines supplement. Culture medium was fully exchanged for medium without CEPT cocktail 24 h after thawing or after each passage and then every 2 d thereafter.

### hiPSC-derived neurons

Excitatory neurons were differentiated from hiPSCs as described before ^63^. hiPSCs were passaged onto Matrigel-treated 35 mm round bottom dishes and transduced on the same day (DIV 0) with a lentivirus expressing the reverse tetracycline transactivator (rtTA) and with a lentivirus expressing Neurogenin-2 (Ngn2) and a puromycin resistance gene under a tetracycline-inducible promoter. After ∼24 h (DIV 1), Ngn2 expression was induced with doxycycline (3 µg/mL) in complete Neurobasal medium composed of Neurobasal medium (Thermofisher A3582901) supplemented with B-27 (Thermofisher A3582801) and Glutamax (Thermofisher 35050061). For selection, puromycin (1 µg/mL) and doxycycline (3 µg/mL) were added to the complete Neurobasal medium for 72 h (DIV 2-4). On DIV5, mouse glial cells were plated on glass coverslips in complete Neurobasal medium supplemented with 5% Fetal Select bovine serum (Atlas Biologicals FS-0500-AD), and induced hiPSCs were recovered from selection in complete Neurobasal medium supplemented with doxycycline (3 µg/mL). Induced hiPSCs were plated on the mouse glial cells ∼24 h later (DIV 6) in complete Neurobasal medium supplemented with doxycycline (3 µg/mL), CEPT cocktail, and 5% Fetal Select bovine serum. Cultures were then maintained until DIV45-55 in complete Neurobasal medium supplemented with 5% Fetal Select bovine serum, human BDNF (10 ng/mL, STEMCELL 78133), human GDNF (10 ng/mL, STEMCELL 78139) and human NT-3 (10 ng/mL, STEMCELL 78074). Half of the culture medium was exchanged every five days. Culture medium was supplemented with doxycycline (3 µg/mL) for DIV7 and 8, as well as with 2-4 μM AraC for DIV7-13 depending on glial growth.

### β4 fusion proteins

All β4 fusion constructs comprise: (1) full-length Ca_V_β4 or truncated derivatives, (2) RIM1α or Munc13-1 sequences, and (3) a C-terminal HA-tag; these components are separated by short, exogenous linkers. Ca_V_β4 and truncated derivatives were generated from a rat Ca_V_β4 cDNA in pMT2 Ca_V_β4 (gift from Annette Dolphin, Addgene Plasmid #107426, lab plasmid code p1251, ^102^), corresponding to Uniprot entry D4A055-2. RIM1 sequences were generated from a rat RIM1 cDNA (p592), corresponding to Uniprot entry Q9JIR4 except for the absence of alternatively spliced exons N_83_IFRC-PERDW_105_, H_1084_SRKS-LLLTRR_1169_, and A_1207_SLVV-QLSQT_1378_ and with two variants (H_263_D and A_453_P; NCBI Reference Sequence NM_052829.3) as described before ^28,36^. Munc13-1 sequences were generated from a rat Munc13-1 cDNA (p194), corresponding to Uniprot entry A0A8I5ZWH4 except for the absence of alternatively spliced exons T_1416_LLRK…SHSDG_1438_ and V_1533_HGGK…VCPEM_1555_. Unless otherwise stated, β4 fusion constructs were cloned into a lentiviral backbone, and expression was driven by the human *SYN1* promoter. The following plasmids were used for this study: pFSW β4-Zn (Lab Plasmid Code p661, described in ^36^), pFSW β4-MC (p1037), pFSW β4-CMC (p1036), pFSW β4-C (p1021), pFSW β4-M (p1020), pFSW β4-M^Δ1Δ2^ (p1055), pFSW β4-M^Δ1Δ2^C (p1056), pFSW β4-M^Δ1^C (p1207), pFSW β4-M^Δ2^C (p1206), pFSW β4-M^Δ1^ (p1205), pFSW β4-M^Δ2^ (p1204), pFSW β4-M^AA^C (p1221), pFSW β4-MC^EE^ (p1222), pFSW β4-M^AA^C^EE^ (p1223), pFSW β4^BID-GK^-Zn (p1218), pFSW β4^GK^-Zn (p1219), pFSW β4^M^-MC (p1232), pFSW β4^S^-MC (p1233), pFSW hSyn^DIO^-β4^S^-Zn (p1238), pFSW SP^DIO^-β4^S^-Zn (p1239, expression is driven by a small promoter, SP, comprised of CMV and human insulin promoter fragments; the short promoter was synthesized based on ^65^ and sequence information available on Addgene, plasmid # 206926; RRID: Addgene_206926), pFSW SP^DIO^-β4^S^-MC (p1240), pAAV hSyn^DIO^-β4^S^-Zn (p1237), pAAV SP^DIO^-β4^S^-MC (p1236). AlphaFold 3 was used to generate 3D models of protein structure ^103^. Models were colorized, annotated and rendered in ChimeraX ^104^.

### Lentivirus

Lentiviruses used to transduce primary mouse hippocampal neurons, hiPSCs and hiPSC-derived neurons were produced in HEK293T cells. HEK293T cells were transfected using the Ca^2+^ phosphate method with viral packaging plasmids REV (p023), RRE (p024) and VSVG (p025), as well as a lentiviral vector encoding the protein of interest. The following lentiviral vectors were used: EGFP-tagged Cre recombinase (to generate cKO^R+E+M^ and cKO^R+E^ neurons, p009); a truncated, enzymatically inactive EGFP-tagged Cre (EGFP-ΔCre, to generate control^R+E+M^ and control^R+E^ neurons, p010); a non-coding vector (to transduce control^R+E+M^, control^R+E^, mouse wild-type, and hiPSC-derived wild-type neurons in place of an expression virus, p008); rtTA (to transduce hiPSCs for Ngn2 induction, Addgene plasmid #20342, gift from R. Jaenisch, p1249 ^105^); Ngn2 (to transduce hiPSCs, Addgene plasmid #52047, gift from M. Wernig, p1250 ^63^); engineered β4 proteins are described above. Plasmids were transfected at a 1:1:1:1 molar ratio. Approximately 24 h after transfection, transfected HEK293T cells were switched into neuronal growth medium (for use on mouse primary neurons) or into hiPSC growth medium (for use on hiPSCs or hiPSC-derived neurons). HEK293T cell supernatant was harvested 36-48 h after transfection following centrifugation for 5 min at 700 x g. For expression of EGFP-Cre and EGFP-ΔCre in mouse primary neurons, neurons were transduced by adding HEK293T cell supernatant at DIV5; for expression of engineered β4 proteins or for the non-coding lentiviral vector in mouse primary neurons, DIV1; for expression of rtTA and Ngn2 in hiPSCs, DIV0; for expression of engineered β4 proteins in hiPSC-derived neurons, DIV5.

### STED microscopy

STED microscopy and analyses were conducted as described before ^28,36,37,56,57,59^. Primary mouse hippocampal neurons were cultured on 12 mm diameter glass coverslips. At DIV15-19, they were washed three times in PBS, fixed in 2% PFA with 4% sucrose (in PBS) for 10 min at room temperature, washed three times in PBS, blocked and permeabilized in PBS with 0.1% Triton X-100 and 3% BSA (TBP) for 1 h at room temperature, then washed again three times in PBS. Primary antibody staining was performed in TBP for 48 h at 4 °C. The following primary antibodies were used: mouse IgG1 anti-HA (1:500, RRID: AB_2565006, lab antibody code A12), rabbit anti-Ca_V_2.1 (1:500, RRID: AB_2619841, A46), guinea pig anti-PSD-95 (1:500, RRID: AB_2619800, A5), guinea pig anti-PSD-95 (1:500, RRID: AB_2905564, A318), mouse IgG1 anti-gephyrin (1:500, RRID: AB_2232546, A8), rabbit anti-gephyrin (1:500, RRID: AB_2619834, A317), rabbit anti-synapsin (1:500, RRID: AB_2200097, A30), mouse IgG1 anti-synapsin (1:500, RRID: AB_2617071, A57), and guinea pig anti-synapsin (1:500, RRID: AB_2905565, A338). Following three washes in PBS, secondary antibody staining was performed in TBP for 24 h at 4 °C. The following secondary antibodies were used: goat anti-mouse IgG1 Alexa Fluor 488 (1:500, RRID: AB_2535764, lab antibody code S7), goat anti-rabbit Alexa Fluor 488 (1:500, RRID: AB_2576217, S5), goat anti-guinea pig Alexa Fluor 555 (1:500, RRID: AB_2535856, S23), goat anti-mouse IgG1 Alexa Fluor 555 (1:500, RRID: AB_2535769, S19), goat anti-rabbit Alexa Fluor 555 (1:500, RRID: AB_2535849, S22), goat anti-rabbit Alexa Fluor 633 (1:500, RRID: AB_2535731, S33), goat anti-mouse IgG1 Alexa Fluor 633 (1:500, RRID: AB_2535768, S29) and goat anti-guinea pig Alexa Fluor 633 (1:500, RRID: AB_2535757, S34). Coverslips were washed twice in PBS and once in deionized water, air-dried and mounted on glass slides with fluorescent mounting medium. STED images were areas of 58.14 x 58.14 μm^2^ (4096 x 4096 pixel^2^, pixel size of 14.194 x 14.194 nm^2^) acquired at 2x digital zoom using a Leica SP8 Confocal/STED 3X microscope equipped with a 100x oil-immersion 1.44 numerical aperture objective, a white light laser, 592 nm, 660 nm and 770 nm depletion lasers, and gated detectors as described before ^59^. Alexa Fluor 633, Alexa Fluor 555, and Alexa Fluor 488 were excited at 633 nm, 555 nm and 488 nm at 1–10% of 1.5 mW laser power. The Alexa Fluor 633 channel was acquired first in confocal mode, then the Alexa Fluor 555 and Alexa Fluor 488 channels were acquired in STED mode using 660 nm and 592 nm depletion lasers, respectively. Identical imaging and laser settings were applied to all conditions within a given biological repeat. To quantify presynaptic localization of HA-tagged β4 fusion proteins or Ca_V_2.1, side-view synapses defined by the presence of a synaptic vesicle cloud (visualized through synapsin) and an elongated postsynaptic density (visualized through PSD-95 or gephyrin) along the edge of the vesicle cloud were selected. Intensity profiles for the postsynaptic density marker and for the protein of interest were generated using a ∼1 μm long, 200 nm wide rectangular ROI drawn perpendicularly through the center of the postsynaptic density. The PSD-95 or gephyrin signal was smoothened for each synapse using a 5-pixel rolling average, and intensity profiles of individual synapses were aligned to the peak PSD-95 or gephyrin signal and averaged. Peak intensities were analyzed by extracting the maximal value from the line profiles of the protein of interest or the smoothened postsynaptic marker within a 200 nm window around the PSD-95 or gephyrin peaks. Peak intensity values were plotted for each synapse and averaged. Representative images in figures were cropped, rotated with bi-linear interpolation, and then brightness and contrast adjusted to facilitate inspection in figures. Brightness and contrast adjustments were done identically for images within an experiment, but image quantification was performed on raw images without these adjustments. The experimenter was blind to the genotype throughout data acquisition and analyses.

### Electrophysiology

Whole-cell recordings were performed similar to methods described before in primary mouse neurons at DIV16-19 ^28,36,59^ and in hiPSC-derived neurons at DIV50-55 ^63,106,107^. Cells were recorded at room temperature in extracellular solution containing (in mM): 140 NaCl, 5 KCl, 2 MgCl_2_, 1.5 CaCl_2_, 10 glucose, and 10 HEPES-NaOH (pH 7.4, ∼300 mOsm). The following drugs were added to the extracellular solution: for electrically evoked NMDAR-EPSCs, 50 μM picrotoxin (PTX) and 20 μM 6-Cyano-7-nitroquinoxaline-2,3-dione (CNQX); for electrically evoked IPSCs, 50 μM D-2-amino-5-phosphonopentanoic acid (D-AP5) and 20 μM CNQX; for electrically evoked AMPAR-EPSCs, 50 μM PTX and 50 μM D-AP5; for sucrose-evoked IPSCs, 1 μM TTX, 20 μM CNQX, and 50 μM D-AP5; for sucrose-evoked AMPAR-EPSCs, 1 μM TTX, 50 μM PTX, and 50 μM D-AP5. Glass pipettes (resistance of 2-5 MΩ) were filled with intracellular solution containing (in mM): for electrically evoked NMDAR- and AMPAR-EPSCs, and for sucrose-evoked AMPAR-EPSCs, 120 Cs-methane-sulfonate, 2 MgCl_2_, 10 EGTA, 4 Na_2_-ATP, 1 Na-GTP, 4 QX314-Cl, 10 HEPES-CsOH (pH 7.4, ∼300 mOsm); and for electrically and sucrose-evoked IPSCs: 40 CsCl, 90 K-gluconate, 1.8 NaCl, 1.7 MgCl_2_, 3.5 KCl, 0.05 EGTA, 2 Mg-ATP, 0.4 Na_2_-GTP, 10 phosphocreatine, 4 QX314-Cl, 10 HEPES-CsOH (pH 7.2, ∼300 mOsm). Cells were held at +40 mV for NMDAR-EPSCs and at −70 mV for IPSCs, and AMPAR-EPSCs. For recordings, access resistance was compensated to 2-3 MΩ, and cells were discarded if the uncompensated access exceeded 15 MΩ at any time point during the recording. For electrically evoked currents, action potentials were evoked with a bipolar focal stimulation electrode made from nichrome wire. For sucrose-evoked currents, 500 mM sucrose in extracellular solution was locally puffed via a microinjector syringe pump for 10 s at a rate of 10 µL/min. Sucrose-evoked currents were quantified as the total charge transfer over the first 10 s following current onset. For paired-pulse ratios, pairs of IPSCs were electrically evoked at interstimulus intervals of 20, 50, 100, 500 and 2500 ms. Paired-pulse ratios were calculated as the peak amplitude of the second postsynaptic current divided by the peak amplitude of the first and were averaged across four trials per interval for each cell. The baseline value for the second postsynaptic current used to derive the peak amplitude was set at the value immediately following the second stimulus artifact. Data were acquired at 10 kHz and lowpass filtered at 2 kHz with an Axon 700B Multiclamp amplifier and digitized with a Digidata 1440A digitizer. Data acquisition and analyses were done using pClamp10. The experimenter was blind to the genotype throughout data acquisition and analyses.

### Electron microscopy

At DIV16–18, neurons cultured on 6 mm sapphire coverslips were high-pressure frozen using a Leica EM ICE high-pressure freezer as described before ^28,36,56,59^. The freezing solution contained: 140 mM NaCl, 5 mM KCl, 2 mM CaCl₂, 2 mM MgCl₂, 10 mM glucose, 10 mM HEPES-NaOH, 50 µM PTX, 20 µM CNQX, and 50 µM D-AP5 (pH 7.4, ∼300 mOsm). Following high-pressure freezing, samples were freeze-substituted in a Leica EM AFS2 system using anhydrous acetone containing 1% glutaraldehyde, 1% osmium tetroxide, and 1% H_2_O. Freeze substitution was performed according to the following temperature program: 5 h at −90 °C, warming from −90 °C to −20 °C at 5 °C/h, 12 h at −20 °C, and warming from −20 °C to 0 °C at 10 °C/h. Samples were washed three times on ice in fresh acetone, three times on ice in propylene oxide, and infiltrated overnight at 4 °C with Epon resin (50% Epon 818 and 50% propylene oxide). The next day, samples were washed with Epon resin three times for 2 h at room temperature, embedded in a mold with fresh resin, and the resin was polymerized at 60 °C for 48 h. The sapphire coverslip was detached from the sample resin by repeated, alternating immersion in liquid nitrogen and warm water. The sample was then sectioned into 50 nm sections using a Leica EM UC7 ultramicrotome, and sections were collected on nickel slot grids (2 x 1 mm) coated with a carbon-supported Formvar film. Sections were counterstained by incubation in 2% lead acetate for 10 s, then rinsed with distilled water. Images were acquired using a JEOL JEM-120i transmission electron microscope at an accelerating voltage of 120 kV and were analyzed using a custom MATLAB script. For each synaptic profile, the number of docked vesicles, the presynaptic bouton area, the number of individual synaptic vesicles, and postsynaptic density length were quantified. Docked vesicles were defined as vesicles for which the electron densities of the vesicular membrane and presynaptic plasma membrane were adjacent, with no visible space between the two membranes. Presynaptic bouton area was determined by tracing the perimeter of each bouton. Active zone length was determined using the electron density of the apposing postsynaptic density or of protein material within the synaptic cleft. The experimenter was blind to the genotype throughout data acquisition and analyses.

### Western blotting

Western blotting was performed as described before ^28,36,56,59^. For western blotting of HEK293T cell homogenates, HEK293T cells were transfected using the Ca^2+^ phosphate method with plasmids expressing the protein of interest and harvested approximately 48 h after transfection. For western blotting of primary mouse neuron and hiPSC-derived neuron homogenates, neurons were transduced with lentiviruses expressing the proteins of interest and harvested at DIV15-19 or DIV50-55, respectively. Homogenates were harvested in standard 3x SDS buffer diluted in PBS, boiled at 95 °C for 10 min, and then centrifuged at 16,200 x g for 10 min. The supernatants were run on 7.5% polyacrylamide gels and transferred onto nitrocellulose membranes for 6.5 h at 4 °C in transfer buffer containing (per L): 200 mL methanol, 14 g glycine and 3 g Tris. Membranes were blocked in filtered 10% nonfat milk/5% goat serum in Tris-buffered saline with 0.1% Tween (TBST) for 1 h at room temperature. Primary antibody staining was performed overnight at 4 °C in 5% nonfat milk/2.5% goat serum in TBST. The following primary antibodies were used: mouse IgG1 anti-HA (1:2500; RRID: AB_2565006, A12), rabbit anti-SNAP-25 (1:10,000; RRID: AB_887790, A151), mouse IgG1 anti-β-actin (1:5000; RRID: AB_476692, A127), rabbit anti-Munc13-1 (1:1000, RRID: AB_887733, A72), rabbit anti-RIM1 (1:1000, RRID: AB_887774, A58) and rabbit anti-synapsin (1:5000, RRID: AB_2200097, A30). Membranes were washed five times in TBST at room temperature, then secondary antibody staining was performed for 1 h at room temperature in 5% nonfat milk/2.5% goat serum in TBST. The following secondary antibodies were used: peroxidase-conjugated goat anti-mouse IgG (1:10,000, RRID: AB_2334540, S52) and peroxidase-conjugated goat anti-rabbit IgG (1:10,000, RRID: AB_2334589, S53). Membranes were again washed five times in TBST at room temperature, incubated in a chemiluminescent reagent for 30 s, and exposed to film that was then developed. Films were scanned as RGB images, and the blue channel converted to grayscale was used for illustration in figures. Images were then rotated with bilinear interpolation and cropped for display.

### Stereotaxic surgeries

Stereotaxic surgeries were performed as described before ^73–75^. Mice were anesthetized with 5% isoflurane and mounted on a stereotaxic frame, and 1.5 to 2% isoflurane was used to maintain stable anesthesia. The scalp was cut to expose the skull, and a small hole was drilled into the skull for syringe insertion. For dopamine sensor imaging, AAV9 expressing GRAB_DA3m_ (BioHippo BHV12400541 ^76^) was injected at a dilution of 1 x 10^12^ copies/mL unilaterally into the right dorsal striatum (coordinates relative to bregma: 1.00 mm anterior, 2.00 mm lateral, 2.50 mm below the dura). For expression of engineered β4 proteins in dopamine neurons, AAV9 (generated by the Boston Children’s Hospital Viral Core from plasmids p1237 or p1236) was injected unilaterally into the right substantia nigra pars compacta (SNc, coordinates relative to bregma: 2.95 mm posterior, 1.25 mm lateral, 4.5 mm below the dura). The following dilutions were used: for AAV9 hSyn^DIO^-β4^S^-Zn, 1.8 x 10^12^ copies/mL; and for AAV9 SP^DIO^-β4^S^-MC, 3.5 x 10^12^ copies/mL. Microinjectors were left in place for 5-10 minutes after completion of injections to allow the injected volume to settle. Analgesia was provided according to approved protocols, and mice were used for further experiments four to five weeks after surgery.

### Slice imaging and analyses

Imaging in acute brain slices was performed as described before ^73,75,108^. Briefly, mice were deeply anesthetized with isoflurane and decapitated. The brain was dissected out, and 250 μm thick sagittal slices were cut using a Leica VT1200S vibratome in ice-cold cutting solution containing (in mM): 75 NaCl, 2.5 KCl, 7.5 MgSO_4_, 75 Sucrose, 1 NaH_2_PO_4_, 12 Glucose, 26.2 NaHCO_3_, 1 Myo-inositol, 3 Na Pyruvic acid, and 1 Na Ascorbic acid. Slices were incubated for 30 min at 37 °C and for 30 min again at room temperature in a recovery solution containing (in mM): 126 NaCl, 2.5 KCl, 2 CaCl_2_, 1.3 MgSO_4_, 1 NaH_2_PO_4_, 12 glucose, and 26.2 NaHCO_3_. Imaging was performed in a chamber continuously perfused at 33-35 °C with a flow rate of 2-3 mL/min of artificial cerebral spinal fluid (ACSF) containing (in mM): 126 NaCl, 2.5 KCl, 2 CaCl_2_, 1.3 MgSO_4_, 1 NaH_2_PO_4_, 12 glucose, and 26.2 NaHCO_3_. For all recordings, ACSF was supplemented with 1 µM dihydro-β-erythroidine (DHβE) to block β2-containing nicotinic receptors. Solutions were constantly bubbled with 95% O_2_ and 5% CO_2_, and data acquisition was completed within 8 hours of slicing. Fluorescence imaging was performed with an Olympus BX51WI epifluorescence microscope. A 470 nm LED was used for excitation, and the measured signals were digitized with a sCMOS camera (Hamamatsu Orca-Flash4.0). A 4x, 0.13 numerical aperture air objective was used for striatal imaging. Electrical stimulation was applied at 90 µA using a pulled glass pipette filled with ACSF (3 to 5 µm tip diameter, 0.5 to 1 MΩ). The following stimulation paradigms were used: single stimulation for two or three replicates, 10-stimulus 10 Hz trains for two or three replicates, or a single 1 Hz train for 130 s (imaging was acquired only for seconds 1-10, 61-70 and 121-130); all replicates were separated by 2-minute interstimulus intervals. A biphasic wave (0.25 ms in each phase) was applied for each stimulus. Fluorescence images were acquired at 512 x 512 pixels/frame and 50 frames/s with an exposure time of 20 ms. Image analyses were performed in Python. For quantification of transients in the dorsal striatum, circular regions of interest (ROIs) with 100-pixel radii were manually selected in each image time series to calculate time series of ΔF/F_0_. The position of each ROI was centered around the visually identified peak signal. To determine baseline fluorescence (F_0_) for fluorescence time series in response to single stimuli or 10 Hz 10 stimuli trains, 200 ms windows before the onset of electrical stimulation were averaged within a ROI. To determine baseline fluorescence (F_0_) for fluorescence time series in response to 1 Hz trains, a single frame prior to each stimulus onset was used. The fluorescence (F) for each time point represents the average pixel intensities within each ROI. ΔF/F_0_ was then calculated as (F-F_0_)/F_0_ for each time point and averaged to generate the time series plots. Peak ΔF/F_0_ values were then extracted from the time series and plotted. Representative heat plot images were made from a 100 ms time window after stimulation, then smoothened with a Gaussian blur (σ = 1) for display in the figures. Recordings from control and experimental conditions were processed identically. The experimenter was blind to the genotype throughout data acquisition and analyses.

### Slice immunofluorescence

Morphological analyses were adapted from protocols used previously ^71,75^. Mice were deeply anesthetized with 5% isoflurane and transcardially perfused with ∼10 mL of phosphate-buffered saline (PBS) followed by ∼10 mL of 4% paraformaldehyde (PFA) in PBS. Brains were post-fixed in 4% PFA in PBS for 12 to 24 h at 4 °C, then stored in PBS at 4 °C. Brains were sectioned into 50 µm slices in PBS using a Leica VT1200S vibratome. Slices were then permeabilized and blocked in PBS containing 0.25% Triton X-100 and 10% normal goat serum for 2 h at room temperature. Primary antibody staining was performed for 24 h at 4 °C in blocking solution, and the following primary antibodies were used: rabbit anti-tyrosine hydroxylase (TH, 1:1000, RRID: AB_390204, A66), mouse IgG1 anti-TH (1:1000, RRID: AB_2201528, A233), mouse IgG1 anti-HA (1:500, RRID: AB_10691311, A250), rabbit anti-HA (1:500, RRID: AB_10693385, A40). Slices were washed three times for 10 min at room temperature in PBS, and secondary antibody staining was performed for 2 h at room temperature in blocking solution. The following secondary antibodies were used: goat anti-mouse IgG1 Alexa Fluor 488 (1:500, RRID: AB_2535764, S7), goat anti-rabbit Alexa Fluor 488 (1:500, RRID: AB_2576217, S5), goat anti-mouse IgG1 Alexa Fluor 555 (1:500, RRID: AB_2535769, S19) and goat anti-rabbit Alexa Fluor 555 (1:500, RRID: AB_2535849, S22). Slices were washed again three times for 10 min at room temperature in PBS and mounted on glass slides for imaging. Confocal images of dorsal striatum were captured on a Nikon ECLIPSE Ti2 inverted spinning disk confocal microscope with a 100x, 0.8 numerical aperture oil-immersion objective. Whole section images containing striatum were acquired using a Leica VS200 brightfield slide scanning microscope with a 10x, 0.6 numerical aperture air objective. Identical image acquisition settings were used for an entire experiment. For quantification of HA levels in dopamine axons labeled by TH, images were median filtered, and background signal was estimated by morphological opening then subtracted for both HA and TH channels. The TH channel only was masked by Otsu thresholding, and the average HA pixel intensity (after filtering and background subtraction) was calculated within this mask. Images were brightness and contrast adjusted to facilitate inspection in figures. These adjustments were done identically for images within an experiment.

## Quantification and statistical analyses

Data are shown as mean ± SEM unless noted otherwise. Statistics were performed in GraphPad Prism 10, and significance is presented as *p < 0.05, **p < 0.01, and ***p < 0.001. Sample sizes and statistical tests for each experiment are included in each figure legend. For STED microscopy in cultured hippocampal neurons, the sample size used for statistical analyses was the number of synapses. For quantifications of bouton area and total vesicles in electron microscopy, the sample size used for statistical analyses was the number of synapses. For quantifications of postsynaptic density length or docked vesicles in electron microscopy, the sample size used for statistical analyses was the number of active zones. For electrophysiological measurements, the sample size used for statistical analyses was the number of recorded cells. For fluorescent sensor imaging in brain slices, the sample size used for statistical analyses was the number of slices. For confocal microscopy of brain slices, the sample size used for statistical analyses was the number of images. Mann-Whitney tests were used for two-condition comparisons. Kruskal-Wallis tests were used for single-factor, multiple group comparisons followed by Dunn’s multiple comparisons post hoc tests. Two-way ANOVA followed by Dunnett’s multiple comparisons tests were used for two-factor, multiple group comparisons. The specific tests used are noted in each figure legend.

---

**Figure S1.**
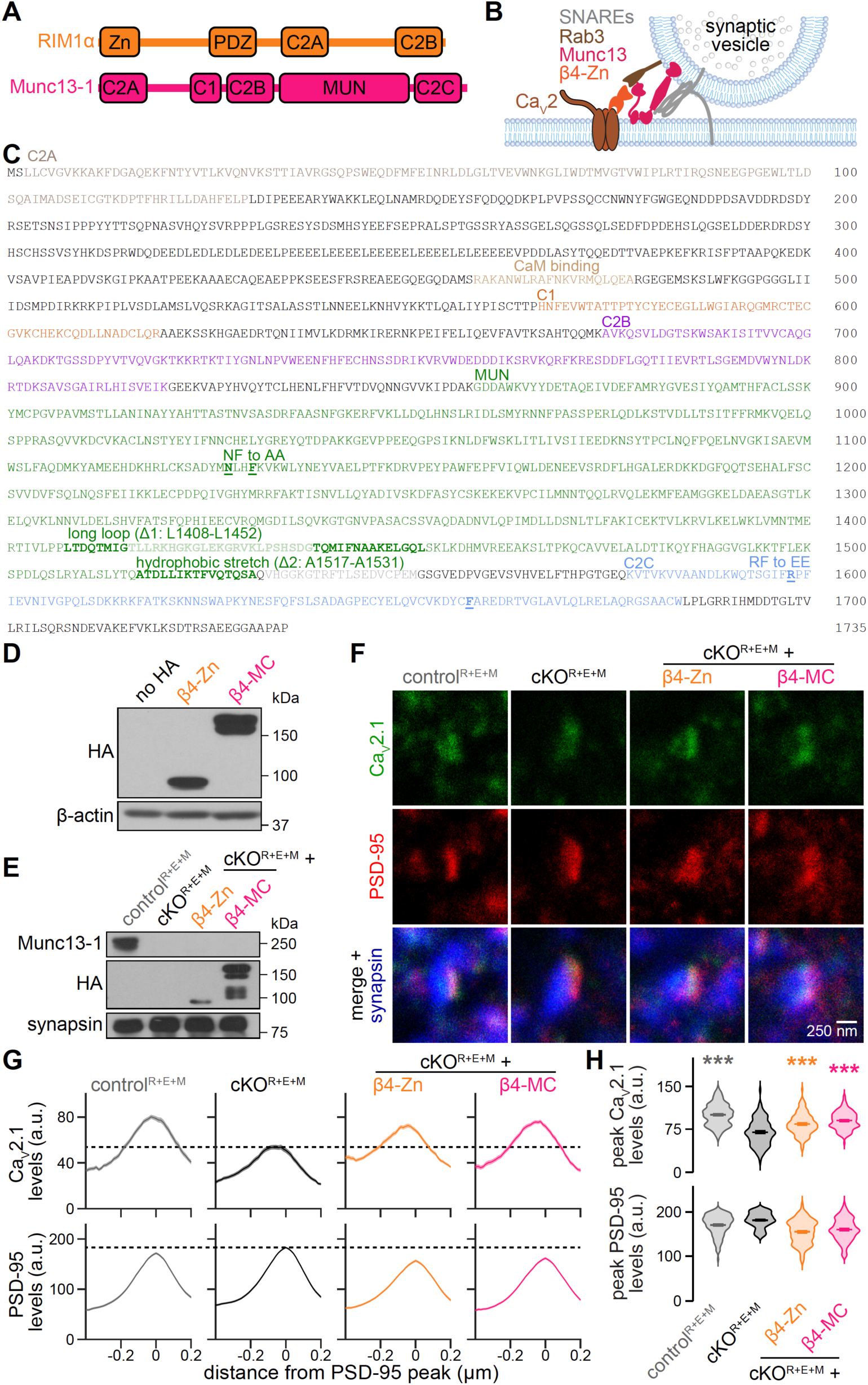
Munc13-1 sequence and additional assessment of β4-Zn and β4-MC. (A) Schematics of RIM1α and Munc13-1, which were used to derive β4-Zn and β4-MC, respectively. (B) Model of β4-Zn function in RIM and ELKS knockout neurons ^36^. (C) Amino acid sequence of rat Munc13-1 (corresponding to Uniprot A0A8I5ZWH4) with annotations of domains, features and tested deletions and point mutations. The sequence of the Munc13-1 cDNA used to generate β4 fusion proteins is identical, except for the absence of residues T_1416_LLRK…SHSDG_1438_ and V_1533_HGGK…VCPEM_1555_ (transparent residues). (D) Western blot of HEK293T cell homogenates without transfection (no HA) or following transfection of β4-Zn or β4-MC; a representative blot from three repeats is shown. (E) Western blot for HA, synapsin and Munc13-1 from homogenates of control^R+E+M^ and cKO^R+E+M^ neurons, and of cKO^R+E+M^ neurons following lentivirally mediated expression of β4-Zn or β4-MC; a representative blot from two repeats is shown. (F-H) Representative images (F) and summary plots of intensity profiles (G) and peak levels (H) of antibody signals at excitatory side-view synapses stained for Ca_V_2.1 (STED), PSD-95 (STED) and synapsin (confocal). Dashed lines in G denote Ca_V_2.1 or PSD-95 levels in cKO^R+E+M^neurons; control^R+E+M^ 133 synapses from 3 cultures, cKO^R+E+M^ 116/3, cKO^R+E+M^ + β4-Zn 125/3, cKO^R+E+M^ + β4-MC 124/3. Data are mean ± SEM; \*\*\**p* < 0.001; statistical significance compared to cKO^R+E+M^ was determined with Kruskal-Wallis tests followed by Dunn’s multiple comparisons post hoc tests in H.

**Figure S2.**
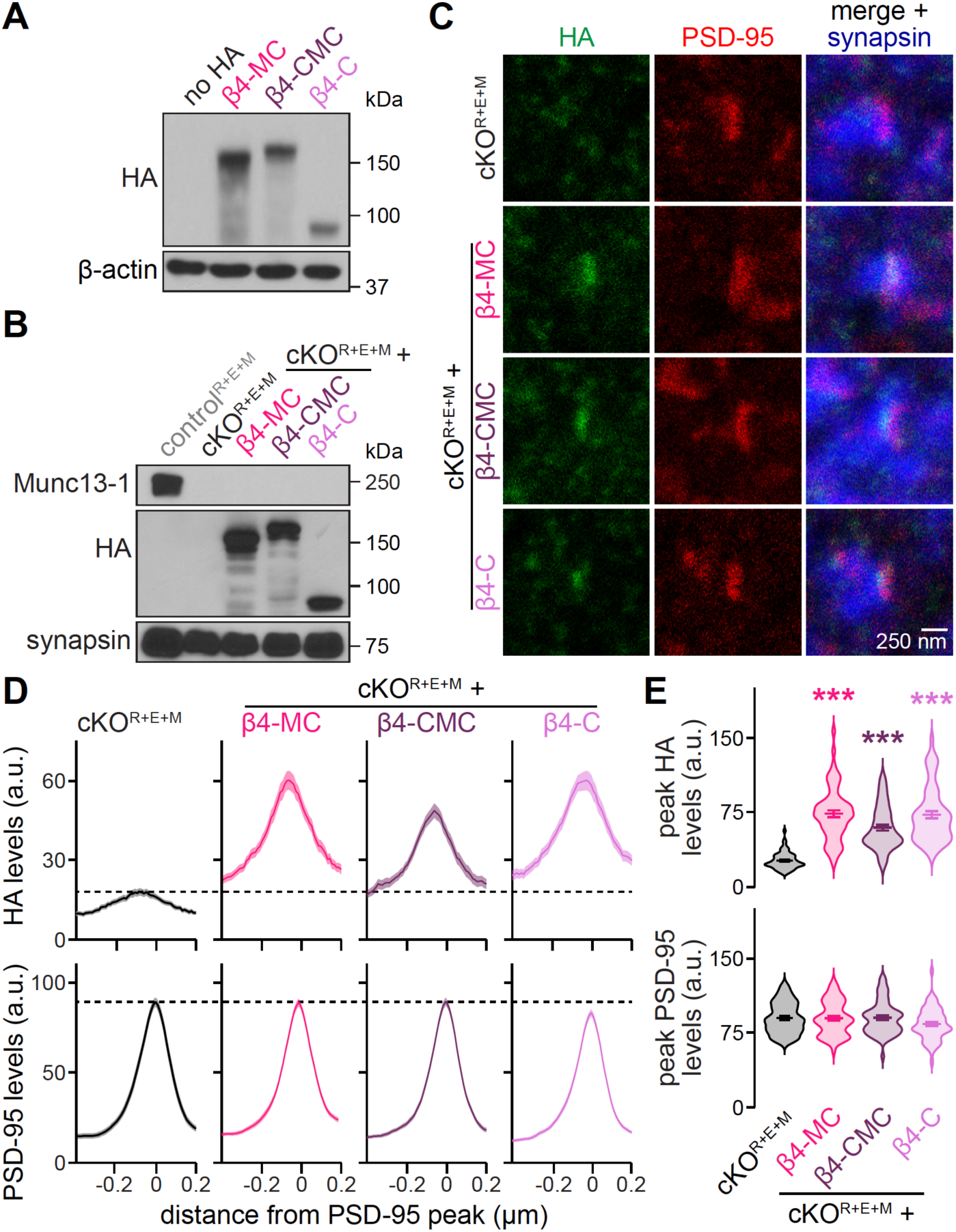
Additional assessment of β4-CMC and β4-C. (A) Western blot for HA and β-actin in HEK293T cell homogenates without transfection (no HA) or following transfection of β4-MC, β4-CMC or β4-C; a representative blot from four repeats is shown. (B) Western blot for HA, synapsin and Munc13-1 from homogenates of control^R+E+M^ and cKO^R+E+M^ neurons, and of cKO^R+E+M^ neurons following lentivirally mediated expression of β4-MC, β4-CMC or β4-C; a representative blot from three repeats is shown. (C-E) Representative images (C) and summary plots of intensity profiles (D) and peak levels (E) of antibody signals at excitatory side-view synapses stained for HA (STED), PSD-95 (STED) and synapsin (confocal). Dashed lines in D denote HA or PSD-95 levels in cKO^R+E+M^ neurons; 60 synapses from 3 cultures each. Data are mean ± SEM; \*\*\**p* < 0.001; statistical significance compared to cKO^R+E+M^ was determined with Kruskal-Wallis tests followed by Dunn’s multiple comparisons post hoc tests in E.

**Figure S3.**
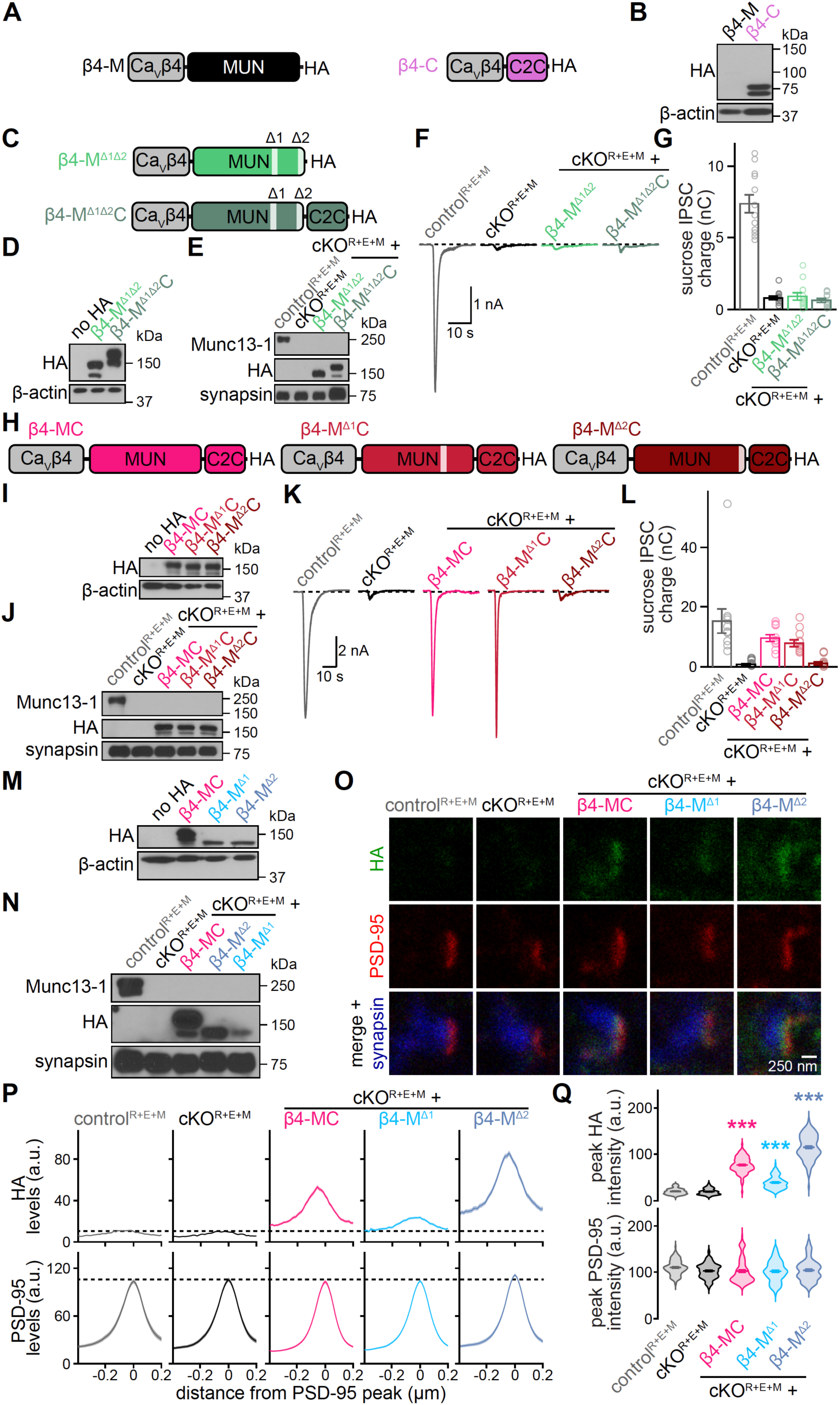
Additional assessment of deletions within the MUN domain. (A) Schematics of β4-M and, for comparison, β4-C. (B) Western blot for HA and β-actin in HEK293T cell homogenates without transfection (no HA) or following transfection of β4-M or, for comparison, β4-C; a representative blot from two repeats is shown. β4-M did not express in transfected HEK293T cells. (C) Schematics of β4-M^Δ1Δ2^ and β4-M^Δ1Δ2^C. (D) Western blot for HA and β-actin in HEK293T cell homogenates without transfection (no HA) or following transfection of the constructs in C; a representative blot from two repeats is shown. Deletions Δ1 and Δ2 enable β4-M^Δ1Δ2^ expression in transfected HEK293T cells. (E) Western blot for HA, synapsin and Munc13-1 from homogenates of control^R+E+M^ and cKO^R+E+M^ neurons, and of cKO^R+E+M^ neurons following lentivirally mediated expression of the constructs in C; a representative blot from two repeats is shown. (F and G) Representative traces (F) and quantification (G) of sucrose-evoked IPSCs recorded from control^R+E+M^ neurons, cKO^R+E+M^ neurons, and from cKO^R+E+M^ neurons expressing the constructs in C; 12 cells from 2 cultures each. Deletions Δ1 and Δ2 abolish β4-M^Δ1Δ2^C function. (H) Schematics of β4-MC, β4-M^Δ1^C and β4-M^Δ2^C. (I) Western blot for HA and β-actin in HEK293T cell homogenates without transfection (no HA) or following transfection of the constructs in H; a representative blot from three repeats is shown.(J) Western blot for HA, synapsin and Munc13-1 from homogenates of control^R+E+M^ and cKO^R+E+M^ neurons, and of cKO^R+E+M^ neurons following lentivirally mediated expression of the constructs in H; a representative blot from two repeats is shown. (K and L) As in F and G but for control^R+E+M^ neurons, cKO^R+E+M^ neurons, and from cKO^R+E+M^ neurons expressing the constructs in H; control^R+E+M^ 11/2, cKO^R+E+M^ 12/2, cKO^R+E+M^ + β4-MC 11/2, cKO^R+E+M^ + β4-M^Δ1^C 11/2, cKO^R+E+M^ + β4-M^Δ2^C 11/2. Restoration of the hydrophobic sequence recovers β4-M^Δ1^C function. (M) Western blot for HA and β-actin in HEK293T cell homogenates without transfection (no HA) or following transfection of β4-MC, β4-M^Δ1^ or β4-M^Δ2^; a representative blot from three repeats is shown. (N) Western blot for HA, synapsin and Munc13-1 from homogenates of control^R+E+M^ and cKO^R+E+M^ neurons, and of cKO^R+E+M^ neurons following lentivirally mediated expression of β4-MC, β4-M^Δ2^ or β4-M^Δ1^; a representative blot from two repeats is shown. (O-Q) Representative images (O) and summary plots of intensity profiles (P) and peak levels (Q) of antibody signals at excitatory side-view synapses of control^R+E+M^ neurons, cKO^R+E+M^ neurons, and cKO^R+E+M^ neurons expressing β4-MC, β4-M^Δ1^ or β4-M^Δ2^. Neurons were stained for HA (STED), PSD-95 (STED) and synapsin (confocal). Dashed lines in P denote HA or PSD-95 levels in cKO^R+E+M^ neurons; control^R+E+M^ 161 synapses/3 cultures, cKO^R+E+M^ 165/3, cKO^R+E+M^ + β4-MC 162/3, cKO^R+E+M^ + β4-M^Δ1^ 169/3, cKO^R+E+M^ + β4-M^Δ2^ 177/3. Data are mean ± SEM; \*\*\**p* < 0.001; statistical significance compared to cKO^R+E+M^ was determined with Kruskal-Wallis tests followed by Dunn’s multiple comparisons post hoc tests in Q. For G and L, no statistical tests were performed because of the limited number of observations.

**Figure S4.**
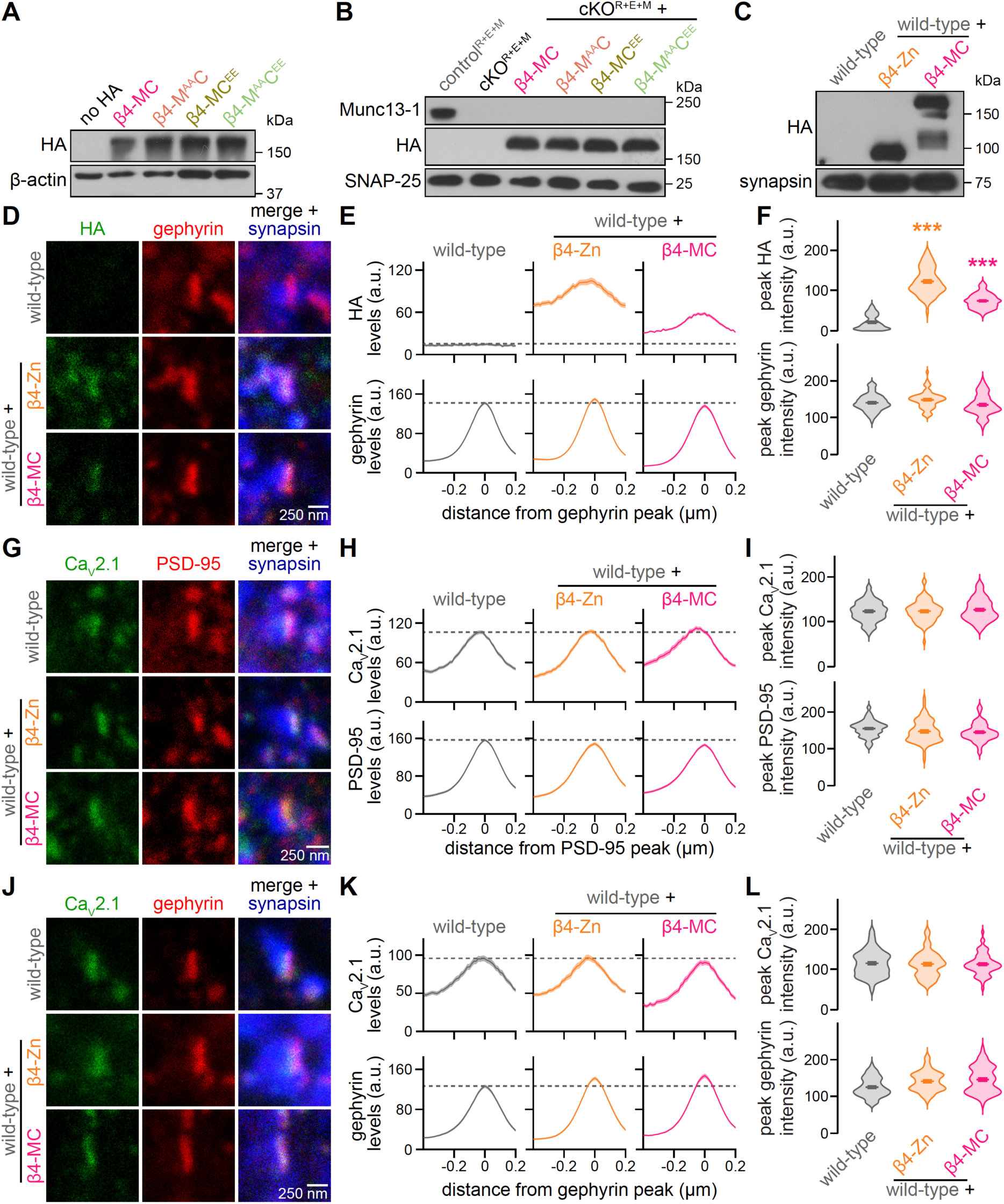
Assessment of β4-M^AA^C, β4-MC^EE^ and β4-M^AA^C^EE^ expression, and of β4-Zn and β4-MC expression and localization in hippocampal wild-type neurons. (A) Western blot for HA and β-actin in HEK293T cell homogenates without transfection (no HA) or following transfection of β4-MC, β4-M^AA^C, β4-MC^EE^, or β4-M^AA^C^EE^; a representative blot from three repeats is shown. (B) Western blot for HA, SNAP-25 and Munc13-1 from homogenates of control^R+E+M^ and cKO^R+E+M^ neurons, and of cKO^R+E+M^ neurons following lentivirally mediated expression of β4-MC, β4-M^AA^C, β4-MC^EE^, or β4-M^AA^C^EE^; a representative blot from five repeats is shown. (C) Western blot for HA and synapsin from homogenates of hippocampal wild-type neurons and of wild-type neurons following lentivirally mediated expression of β4-Zn or β4-MC; a representative blot from six repeats is shown. (D-F) Representative images (D) and summary plots of intensity profiles (E) and peak levels (F) of antibody signals at inhibitory side-view synapses. Neurons were stained for HA (STED), gephyrin (STED) and synapsin (confocal). Dashed lines in E denote HA or gephyrin levels in wild-type neurons; wild-type 70 synapses from 3 cultures, wild-type + β4-Zn 68/3, wild-type + β4-MC 72/3. (G-I) As in D-F but for excitatory synapses stained for Ca_V_2.1, PSD-95 and synapsin; wild-type 80/3 cultures, wild-type + β4-Zn 78/3, wild-type + β4-MC 75/3. (J-L) As in D-F but for inhibitory synapses stained for Ca_V_2.1, gephyrin and synapsin; wild-type 70/3, wild-type + β4-Zn 67/3, wild-type + β4-MC 73/3. Data are mean ± SEM; \*\*\**p* < 0.001. Statistical significance compared to wild-type was determined with Kruskal-Wallis tests followed by Dunn’s multiple comparisons post hoc tests in F.

**Figure S5.**
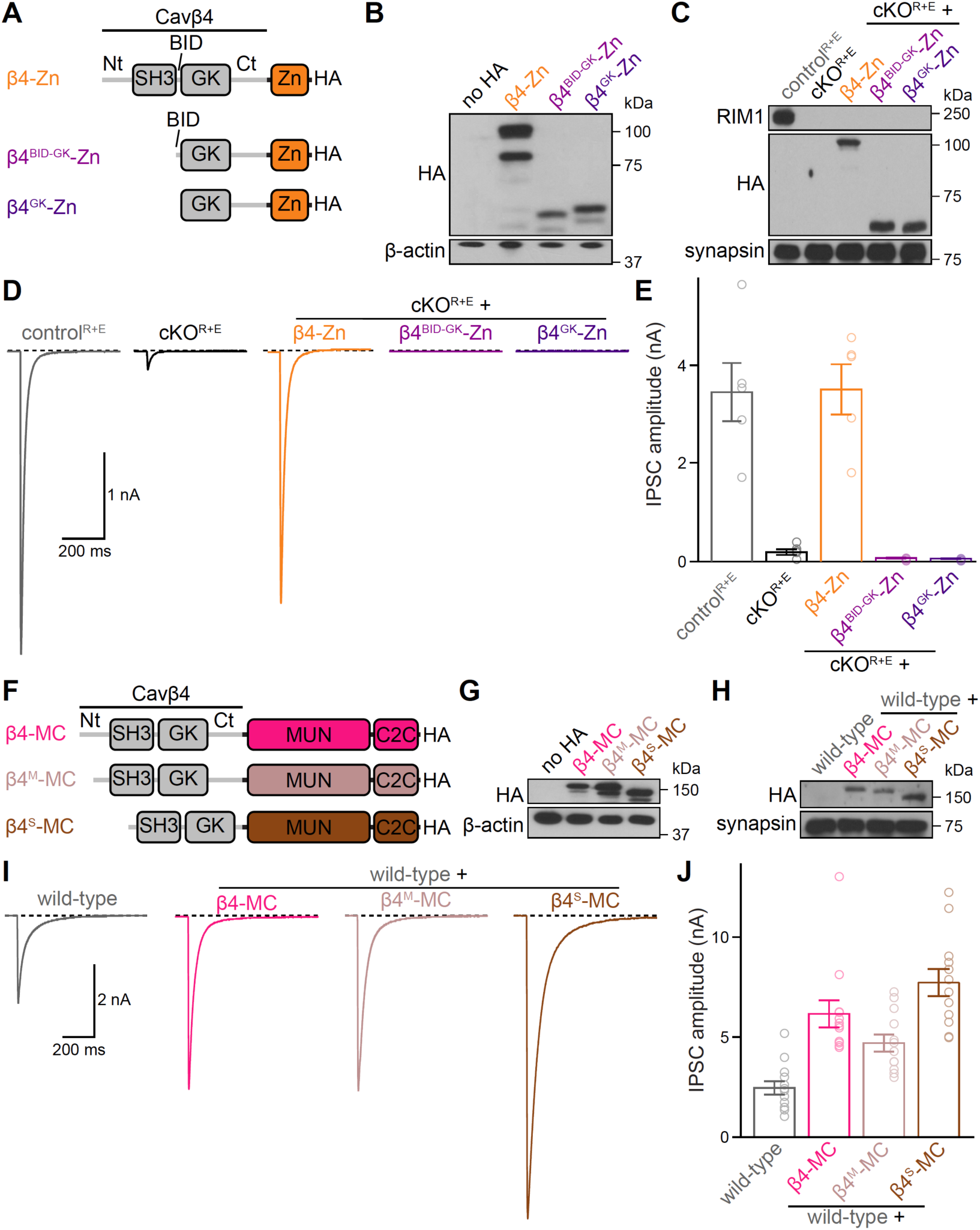
Generation of short Ca_V_β4 sequences for engineered β4 proteins. (A) Schematics of β4-Zn, β4^BID-GK^-Zn and β4^GK^-Zn. (B) Western blot for HA and β-actin in HEK293T cell homogenates without transfection (no HA) or following transfection of the constructs in A; a representative blot from two repeats is shown. (C) Western blot for HA, synapsin and RIM1 from homogenates of control^R+E^ and cKO^R+E^ neurons, and of cKO^R+E^ neurons following lentivirally mediated expression of the constructs in A; a representative blot from two repeats is shown. (D and E) Representative traces (D) and quantification (E) of electrically evoked IPSCs recorded from control^R+E^ and cKO^R+E^ neurons, and from cKO^R+E^ neurons expressing the constructs in A; 5 cells from 1 culture each. (F) Schematics of β4-MC, β4^M^-MC and β4^S^-MC. (G) Western blot for HA and β-actin in HEK293T cell homogenates without transfection (no HA) or following transfection of the constructs in F; a representative blot from two repeats is shown. (H) Western blot for HA and synapsin from homogenates of hippocampal wild-type neurons and of wild-type neurons following lentivirally mediated expression of the constructs in F; a representative blot from three repeats is shown. (I and J) As in D and E but for electrically evoked IPSCs recorded from wild-type neurons, and from wild-type neurons expressing the constructs in F; 12/2 each. Data are mean ± SEM; no statistics were performed in E and J because of the limited number of observations.

**Figure S6.**
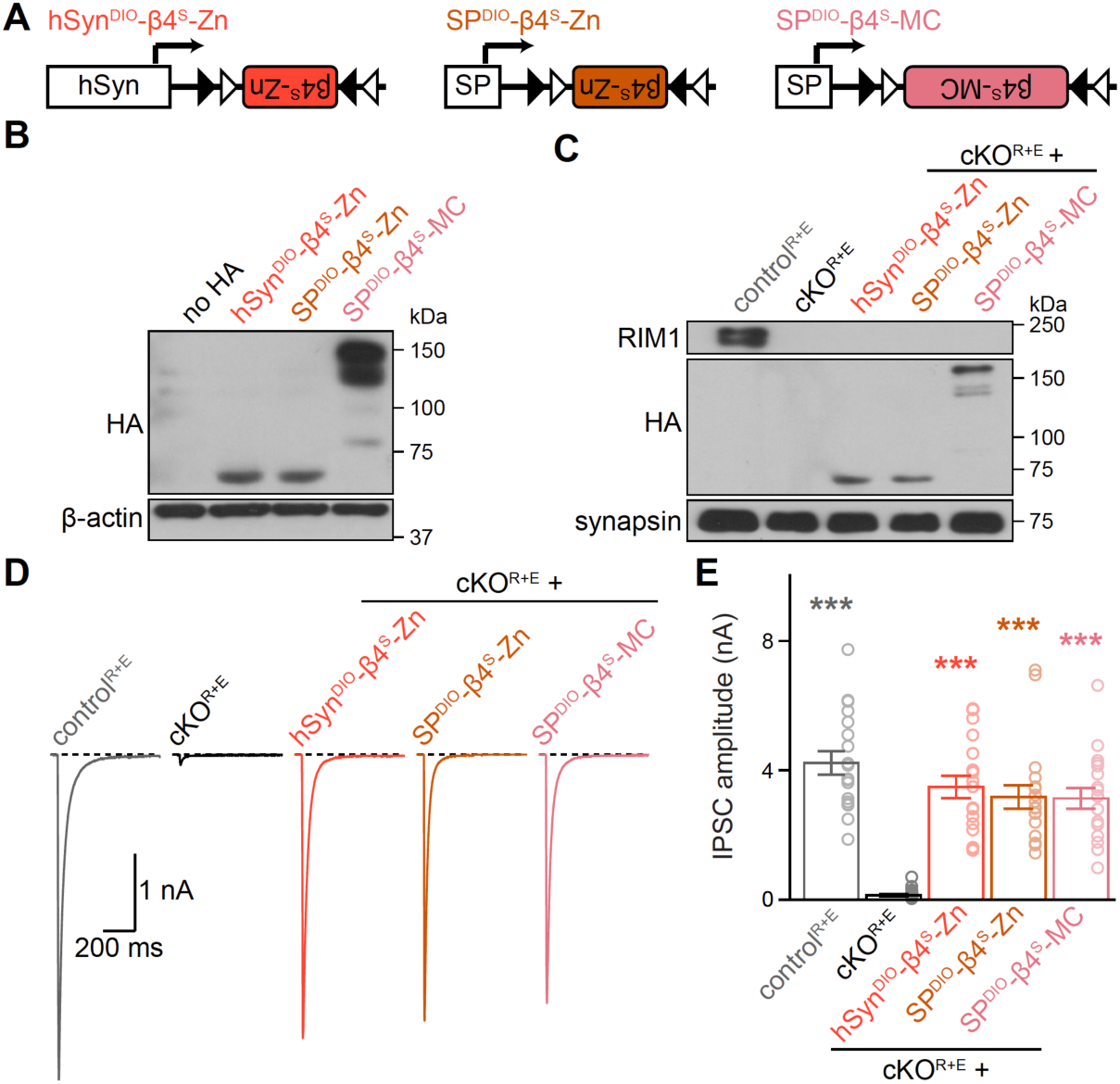
Conditional expression of engineered β4 proteins from a short promoter. (A) Schematics of hSyn^DIO^-β4^S^-Zn, SP^DIO^-β4^S^-Zn and SP^DIO^-β4^S^-MC. For SP^DIO^-β4^S^-Zn and SP^DIO^-β4^S^-MC, expression is driven by a synthetic, short promoter (SP) ^65^. (B) Western blot for HA and β-actin from HEK293T cell homogenates without transfection (no HA) or following transfection of Cre recombinase and the constructs in A; a representative blot from three repeats is shown. (C) Western blot for HA, synapsin and RIM1 from homogenates of control^R+E^ and cKO^R+E^ neurons, and of cKO^R+E^ neurons following lentivirally mediated expression of the constructs in A; a representative blot from two repeats is shown. (D and E) Representative traces (D) and quantification (E) of electrically evoked IPSCs; 18 cells from 3 cultures each. Data are mean ± SEM; \*\*\**p* < 0.001. Statistical significance compared to cKO^R+E^ was determined with Kruskal-Wallis tests followed by Dunn’s multiple comparisons post hoc tests in E.

**Figure S7.**
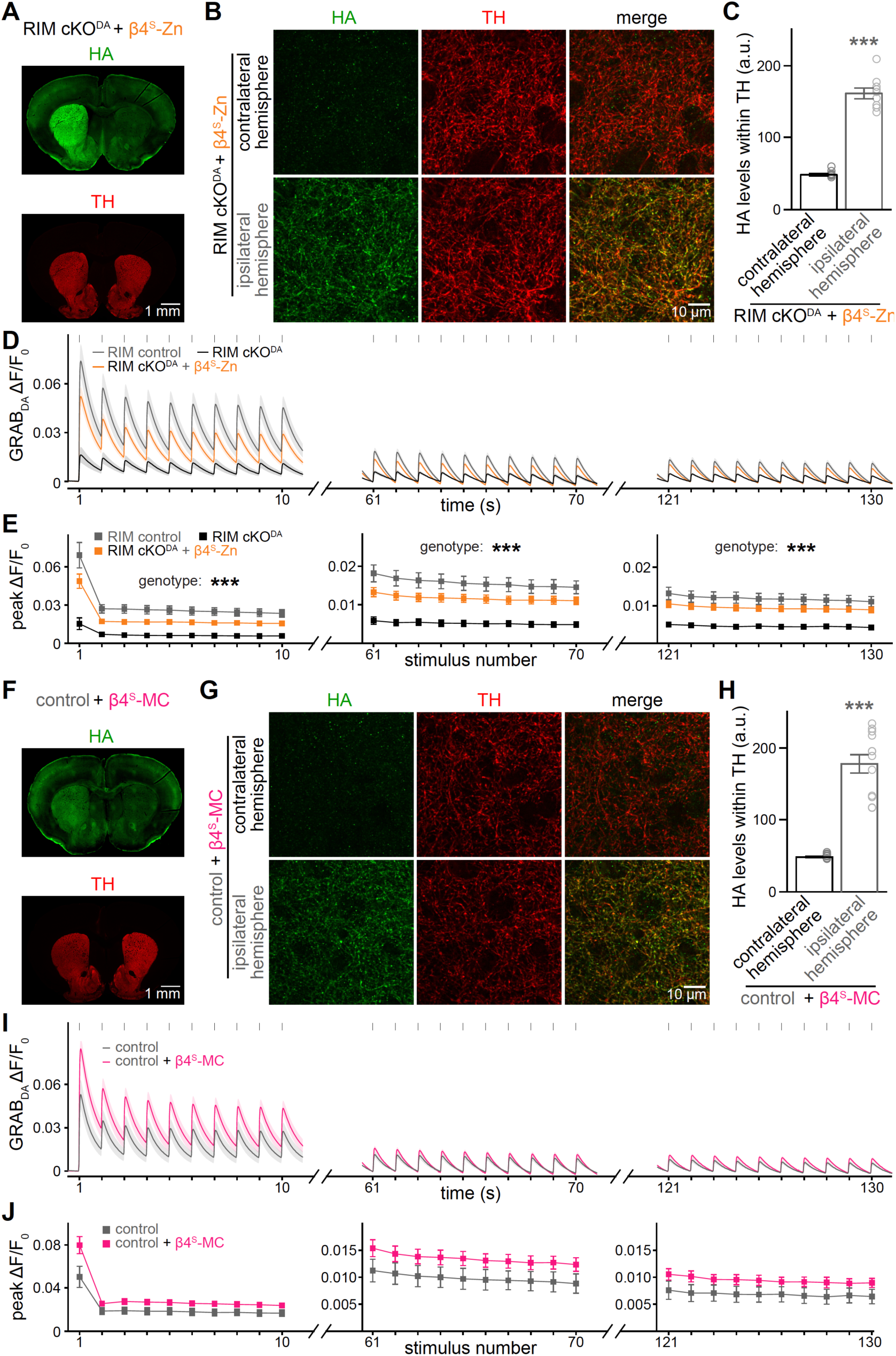
Assessment of β4 protein expression in striatal dopamine axons and of striatal dopamine release in response to 1 Hz stimulation. (A-C) Representative 10x slide scanner images (A), representative 100x confocal images (B) and quantification of HA levels in dopamine axons (C) within the dorsolateral striatum of RIM cKO^DA^ mice following unilateral injections of AAV9-hSyn^DIO^-β4^S^-Zn in the SNc. Coronal sections were stained for HA to visualize β4^S^-Zn and for tyrosine hydroxylase (TH) to visualize dopamine axons; 9 images from 3 mice each. (D and E) Quantification of the GRAB_DA3m_ ΔF/F_0_ time course (D) and the ΔF/F_0_ peak per stimulus (E) during 130 s long 1 Hz stimulation in the dorsolateral striatum. Imaging was only acquired at 1 s to 10 s, 61 s to 70 s, and 121 s to 130 s. Vertical ticks in (D) denote the electrical stimuli; RIM control 11 slices from 4 mice, RIM cKO^DA^ 14/4, RIM cKO^DA^ + β4^S^-Zn 13/4. (F-H) As in A-C but for control mice injected with AAV9-SP^DIO^-β4^S^-MC, all mice had a DAT^IRES-Cre^ allele; 12/3 each. (I and J) As in D and E but for control mice and control mice expressing β4^S^-MC, all mice had a DAT^IRES-Cre^ allele; control 8/3, control + β4^S^-MC 9/3. Data are mean ± SEM. \*\*\**p* < 0.001, statistical significance compared to the contralateral hemisphere was determined with Mann-Whitney tests in C and H. Statistical significance in E was determined by a two-way ANOVA followed by Dunnett’s multiple comparisons tests compared to RIM cKO^DA^. In E, stimulus number 1-10: genotype *p* < 0.001, stimulus number *p* < 0.001, interaction *p* < 0.001; stimulus number 61-70: genotype *p* < 0.001, stimulus number *p* < 0.001, interaction *p* < 0.001; stimulus number 121-130: genotype *p* < 0.001, stimulus number *p* < 0.001, interaction *p* = 0.009; Dunnett’s multiple comparisons post hoc tests were performed for stimulus number 1 (RIM control vs. RIM cKO^DA^ *p* < 0.001, RIM cKO^DA^ + β4^S^-Zn vs. RIM cKO^DA^ *p* < 0.001), stimulus number 10 (RIM control vs. RIM cKO^DA^ *p* < 0.001, RIM cKO^DA^ + β4^S^-Zn vs. RIM cKO^DA^ *p* < 0.001), stimulus number 70 (RIM control vs. RIM cKO^DA^ *p* < 0.001, RIM cKO^DA^ + β4^S^-Zn vs. RIM cKO^DA^ *p* < 0.001), and stimulus number 130 (RIM control vs. RIM cKO^DA^ *p* < 0.001, RIM cKO^DA^ + β4^S^-Zn vs. RIM cKO^DA^ *p* < 0.001).

